# Transcriptional kinetics and molecular functions of long non-coding RNAs

**DOI:** 10.1101/2020.05.05.079251

**Authors:** Per Johnsson, Christoph Ziegenhain, Leonard Hartmanis, Gert-Jan Hendriks, Michael Hagemann-Jensen, Björn Reinius, Rickard Sandberg

## Abstract

An increasing number of studies have demonstrated the regulatory importance of long non-coding RNAs (lncRNAs), yet little is known about their transcriptional dynamics and it remains challenging to determine their regulatory functions. Here, we used allele-sensitive single-cell RNA-seq (scRNA-seq) to demonstrate that lncRNAs have lower burst frequencies with twice as long duration between bursts, compared to mRNAs. Additionally, we observed an increased cell-to-cell variability in lncRNA expression that was due to more sporadic bursting (lower frequency) with larger numbers of RNA molecules being produced. Exploiting heterogeneity in asynchronously growing cells, we identified and experimentally validated lncRNAs with cell-state specific functions involved in cell cycle progression and apoptosis. Finally, utilizing allele-resolved RNA expression, we identified *cis* functioning lncRNAs and observed that knockdown of these lncRNAs modulated either transcriptional burst frequency or size of the nearby protein-coding gene. Collectively, our results identify distinct transcriptional regulation of lncRNAs and we demonstrate a role for lncRNAs in the regulation of transcriptional bursting of mRNAs.

## Introduction

Mammalian genomes encode thousands of lncRNAs^1^, but identifying their molecular functions has proven difficult. Functional predictions based on primary sequence, evolutionary conservation^2^, or genomic location are often unreliable and to date we still cannot identify active lncRNAs and their mechanism of action without extensive experimental validation. Consequently, the functions of most lncRNAs remain unknown^3^ and new experimental and computational approaches are highly needed in order to efficiently identify lncRNAs for in-depth functional validation and characterization.

Numerous studies of lncRNAs have shown that their average expression levels are lower than those of mRNAs^4–9^, yet vital regulatory functions have been demonstrated for several lncRNAs^10–12^. The observed low expression of lncRNAs is generally oversimplified by assuming that all cells in a population have uniform expression, i.e. resulting in expression estimates below one RNA copy per cell^13^ that are not easy to functionally perceive. It has been proposed that averaging transcriptomes over thousands of cells could mask the presence of few cells with relatively high expression of specific lncRNAs^14^. However, comprehensive analyses of transcriptional dynamics and cell-to-cell variability of lncRNAs are still missing, and most studies to date were limited to low throughput methods covering low numbers of genes and cells^15^. With the introduction of scRNA-seq technologies^16^ and specifically protocols allowing for allele-specific quantification^17^, it is now feasible to characterize allele specific gene expression in individual cells for thousands of genes simultaneously. Although scRNA-seq is a powerful tool for identifying cell types^18^, transient cellular states^19^ and burst kinetics^20^, dedicated scRNA-seq studies with allelic resolution focusing on lncRNAs are so far lacking.

Transcription of mammalian genes typically occurs in short bursts of activity^21^, where burst frequencies are generally controlled by enhancers, whereas burst sizes are in part dependent upon core promoter elements^20^. Through recent methodological^22^ and computational^20^ developments, it is now feasible to infer burst parameters for thousands of genes simultaneously. To date, analysis have mainly focused on protein-coding genes and it remains unknown for example whether low expression of lncRNAs is mediated by lowered burst sizes (fewer RNA molecules per cell) or burst frequencies (expression in fewer cells). Recent studies have suggested lncRNA promoters to have fewer transcription factor (TF) motifs^5,23^ and less efficient polymerase II pausing^24^, than those of mRNAs. How these observations effect transcriptional bursts of lncRNAs, and whether lncRNAs have distinct or similar burst characteristics to mRNAs, remains unknown.

In this study, we introduce allele-sensitive scRNA-seq to investigate transcriptional dynamics and molecular functions of lncRNAs. We first set out to study burst kinetics of lncRNAs and investigate if lncRNAs have different burst kinetics compared to mRNAs. We next explore if the identification of transient cellular states with precise expressions of lncRNAs predicts their molecular functions. Using an expression-to-phenotype relationship in non-perturbed asynchronously growing cells, we identify several lncRNAs involved in cell cycle regulation as well as apoptosis. Next, we explore patterns of allelic expression of proximal lncRNA-mRNA gene pairs and functionally confirm that this is a powerful approach to map *cis* functioning lncRNAs. We finally provide insights into how lncRNAs modulate burst dynamics of *cis*-regulated genes and show that lncRNAs can modulate both burst frequencies and burst sizes. In summary, our study provides novel and comprehensive insights into burst dynamics of lncRNAs and demonstrates that allele-sensitive scRNA-seq is a powerful tool for in-depth characterization of lncRNAs.

## Results

### Detection of lncRNAs and mRNAs in individual cells

We first investigated the expression patterns of lncRNAs and mRNAs in individual primary mouse fibroblasts derived from the cross between the distantly related CAST/EiJ and C57BL/6J strains. To this end, we profiled 375 individual adult tail fibroblasts with Smart-seq2 to leverage the methods’ high sensitivity^25^ and full gene body coverage enabling allele-level RNA profiling for more than 80% of all genes^17^. We also made use of a previously published data set consisting of additional 158 cells, generating a comprehensive data set of 533 deep-sequenced fibroblasts (4.2 million mapped reads per cell, median, **Figure S1A**). For allelic expression estimates, the fraction of allele-informative bases in reads covering heterozygous single-nucleotide polymorphisms (SNP) was used. To rule out preferential mapping or misalignment we verified that non-imprinted autosomal genes (**Supplementary Table 1**) had similar overall expression from the CAST and C57 alleles (**Figure S1B**) and that we accurately detected monoallelic expression for non-escapee X-chromosomal genes^26^ (**Figure S1C, Supplementary Table 2**). A total number of 24,653 genes were detected (requiring 5 or more read counts in at least 2 cells), including 15,872 mRNAs and 3,314 non-coding RNAs. The detection of hundreds of lncRNAs in individual cells (9,174 protein-coding mRNAs and 408 lncRNAs per cell, median, **Figure 1A**), motivated us to proceed with in-depth investigations of lncRNAs. Since lncRNAs are a diverse group of transcripts where we noticed an effect on gene expression levels on closely located promoters (**Figure S1D**), we focused our analysis on easily separated transcriptional units (**Figure S1E**).

**Figure 1.**
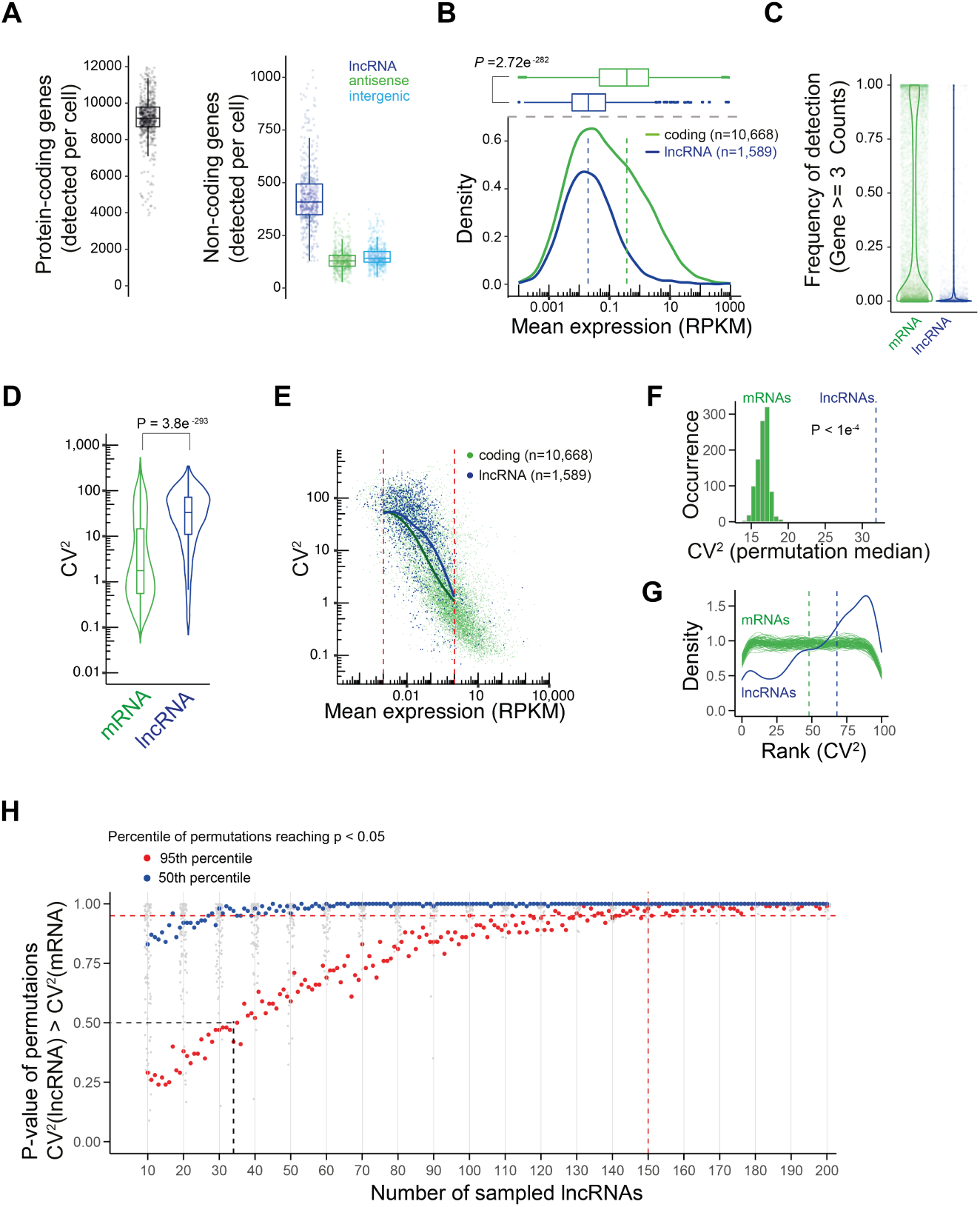
Levels and variability of lncRNA and mRNA expressions. **(A)** Boxplots showing the detected numbers of protein-coding genes and subtypes of lncRNAs per fibroblast, based on Smart-seq2 data (n=533 cells) and requiring 3 or more read counts for detection. **(B)** Densities and boxplots of mean expression levels for lncRNAs and mRNAs across fibroblasts (n=533). Dashed lines denote the medians, p-value represents a two-sided Wilcoxon rank-sum test. **(C)** Violin plots showing the fraction of cells that detected individual lncRNAs and mRNAs (requiring 3 or more read counts for detection). **(D)** Violin plots showing the coefficient of variation (CV^2^) for lncRNAs and mRNAs expressions across fibroblast (n=533). P-value represents two-sided Wilcoxon rank-sum test. **(E)** Scatter plot of mean expression against the CV^2^ for lncRNAs (blue) and mRNAs (green). Lines denote a smoothed fit to the rolling mean (width = 15) for lncRNAs and mRNAs. Red dotted lines denote the expression range for the smoothed fit. **(F)** Histogram showing median CV^2^ for sampled expression-matched sets of mRNAs. Each lncRNA (0.001 < lncRNA_mean_ < 100, n=1,519) was matched with 10 mRNAs of similar expression followed by subsampling one expression-matched mRNA for each lncRNA, this procedure was repeated 10,000 times. The P-value represent the outcome of the permutation test where the CV^2^_mRNA_ obtained from 10,000 permutations was higher than the observed CV_2lncRNA_ (blue dashed line). **(G)** Densities of rankings of CV^2^ for lncRNAs (blue, n=1,519) and randomly sampled mRNAs (green). In blue, each lncRNA was matched with 100 mRNAs of similar expression followed by ranking the CV^2^ to the 100 matched mRNAs (frequency CV^2^_lncRNA_ > CV^2^_mRNA_matched_). In green, mRNAs were randomly sampled (n=1,519, as many as lncRNAs), expression-matched with 100 other mRNAs and the CV^2^ ranked (frequency CV^2^_mRNA_random_ > CV^2^_mRNA_matched_). This procedure was repeated 100 times for mRNAs. Dashed lines denote the medians of ranking for lncRNAs (blue) and mRNAs (green). **(H)** Scatter plot showing the numbers of lncRNAs required to identify increased CV^2^ compared to expression-matched mRNAs. The X-axis represents the number of lncRNAs used for CV^2^ quantification. For each number of lncRNAs analyzed, expression-matched mRNAs were randomly selected followed by subsampling one expression-matched mRNA for each lncRNA (similar as in Figure 1G, 1,000 permutations). Each subsampling of lncRNAs was repeated 100 times. The y-axis represents p-values from the permutation test (median(CV^2^_lncRNA_) > median(CV^2^_mRNA_)). Grey points represent individual p-values for the permutation test. Blue points represent the 50^th^ percentile of permutations reaching significance. Red points represent the 95^th^ percentile of permutations reaching significance. The black dashed line represents subsampling 34 lncRNAs. Analysis in Figures 1D-G represent easily separated transcriptional units of lncRNAs and mRNAs.

### lncRNAs are expressed with higher cell-to-cell variability than mRNAs

We first investigated the patterns of expression of lncRNAs and mRNAs, and as expected^4,27^, we observed that lncRNAs were expressed at lower levels than mRNAs (**Figure 1B**) and detected in fewer cells (median 3% and 31% of cells, respectively) (**Figure 1C**). To further investigate if lncRNAs are expressed with higher variability between cells, we computed the squared coefficient of variation (CV^2^) and observed significantly higher variability for lncRNAs (**Figure 1D**, *P*=3.8e^-293^, two-sided Wilcoxon rank-sum test). Contrasting CV^2^ against the mean expression revealed that lncRNAs had higher CV^2^ than mRNAs across a wide range of expression levels (**Figure 1E**). To systematically account for possible confounding differences in mean expression of lncRNAs and mRNAs, we generated thousands of randomly drawn sets of mRNAs with expressions matched to lncRNAs (**Figure 1F**, see Methods) and also ranked the CV^2^ of each lncRNA against 100 expressions matched mRNAs (**Figure 1G**). Consistently, lncRNAs had significantly higher expression variability than expression-matched mRNAs (P<1e^-4^, permutation test) (**Figures 1F-G**). The increased cell-to-cell variability was also validated in human HEK293 cells (**Figure S2A**), as well as mouse embryonic stem cells (**Figure S2B**). The ability to detect the increased cell-to-cell variability was dependent on the number of lncRNAs analyzed, and when subsampling lncRNAs (and their expression-matched mRNAs) the difference declined and eventually disappeared (**Figure 1H**). The lack of power when analyzing small numbers of lncRNAs might explain why a previous study investigating 34 lncRNAs found no difference in variability compared with mRNAs^15^.

### Low expression of lncRNAs results from longer duration between bursts

We next sought to determine how the lowered expression level of lncRNAs is achieved in terms of transcriptional bursting kinetics and whether lncRNAs are intrinsically different from protein-coding genes. To do this, we generated a comprehensive data set of adult tail fibroblast using Smart-seq3^22^ (682 cells post quality control, 3.0 Million mapped 100bp paired-end reads per cell, median, **Figures S3A-C**) which, in addition to Smart-Seq2, provides unique molecular identifiers (UMIs) important for accurate burst size inference^20^. After quality control of genes (**Figures S3D-E**), bursting kinetics parameters were successfully inferred for 10,716 coding genes and 655 lncRNAs on at least one of the alleles (8,661 coding and 325 lncRNAs genes on both alleles). Reassuringly, burst parameters and expression levels correlated well between the CAST and C57 alleles for both coding and non-coding genes (**Figures 2A-C**). Focusing the analysis on separated transcriptional units (**Figure S1E**) we found that lncRNAs have a four-fold lower burst frequency compared to mRNAs (**Figures 2D** and **S4A**), and only a two-fold decrease in burst size (**Figures 2E** and **S4B**). Thus, the decreased expression of lncRNAs (**Figures 2F** and **S4C**) is mainly achieved through longer duration between bursts of expression, possibly regulated by enhancer activities^20,28–30^.

**Figure 2.**
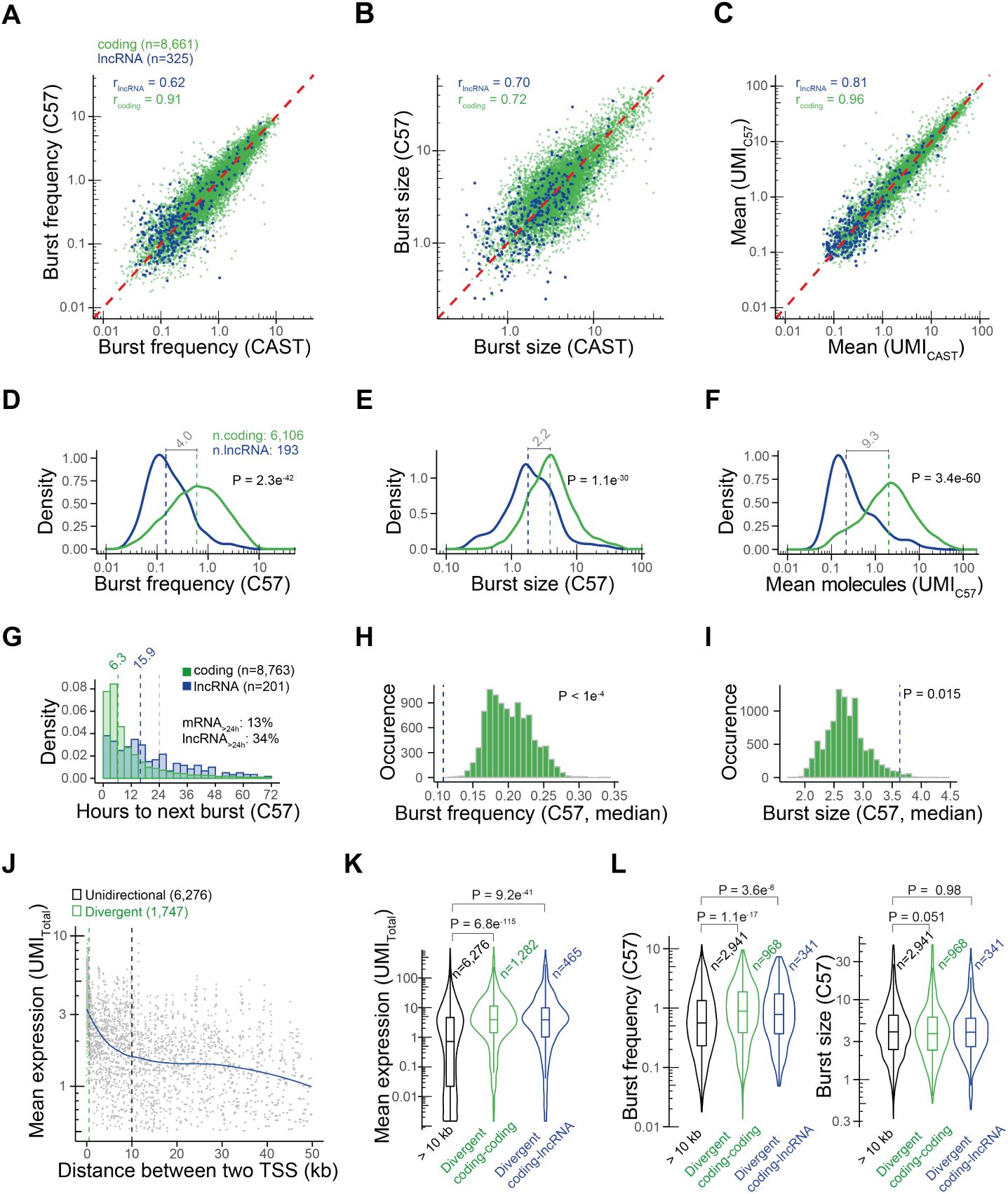
Transcriptional burst kinetics of lncRNAs and divergent promoters. **(A-C)** Scatter plots of (A) burst frequencies, (B) burst sizes and (C) mean expression for mRNAs (green) and lncRNAs (blue) comparing the parameters inferred from the CAST allele against the C57 allele for non-imprinted autosomal genes. Red line denotes *x=y*. **(D-F)** Density plots for (D) burst frequencies (E) burst sizes and (F) mean expression (allele-distributed UMIs) for mRNAs and lncRNAs (showing the C57 allele). Dashed lines represent the median burst frequencies, sizes and mean expression for mRNAs and lncRNAs. The relative fold changes in burst frequencies, sizes and mean expression are annotated in grey. The p-values represent two-sided Wilcoxon rank-sum tests. **(G)** Histogram showing the duration between two bursts from the same allele, for mRNAs and lncRNAs. Dashed lines in green and blue represent the median duration between two bursts for mRNAs and lncRNAs, respectively. Dashed line in grey represents a duration of 24 hours between two bursts. **(H-I)** Histograms showing medianM (H) burst frequencies (I) and burst sizes for sampled expression-matched sets of mRNAs. Each lncRNA (n=50, identified in Figure S4H) was matched with 10 mRNAs of similar expression followed by subsampling one expression-matched mRNA for each lncRNA. The P-value represent the outcome of the permutation test, where the observed burst parameters (lncRNAs, median) was higher (for burst frequencies) or lower (for burst sizes) than the burst parameters (median) obtained from 10,000 permutations. **(J)** Scatter plot showing the distance between the transcription start sites (TSS) of pairs of genes, against their mean expression levels (UMIs). Blue line represents a loess fit to the rolling median (width = 31). The dashed lines represent the distance between two TSSs for being assigned as divergent promoters (in green, maximum distance of 500 bp) or unidirectional promoters (in black, minimum distance of 10 kb). **(K)** Violin plots showing mean expression levels of unidirectional mRNAs and for mRNAs transcribed from divergent promoters (either with another mRNA or a lncRNA). P-values represent two-sided Wilcoxon rank-sum tests. **(L)** Violin plots for unidirectional and divergent promoters representing burst frequencies and burst sizes, for the C57 allele. P-values represent two-sided Wilcoxon rank-sum test.

Since the inferred parameters for burst frequencies are on the time-scale of RNA degradation^20^, we next sought to derive burst frequencies on an absolute time-scale, which requires information of RNA decay rates. To address this, we measured RNA half-lives in primary fibroblasts upon transcriptional inhibition with actinomycin D and fit the normalized expression of each gene to a first order exponential decay curve (see Methods). The estimates were in agreement with previous measurements (**Figure S4D**)^31^, with an average half-life slightly below four hours (**Figure S4E**). Furthermore, our data do not support any systematic difference in half-lives (**Figure S4E**) or decay rates (**Figure S4F**) between mRNAs and lncRNAs, thus in line with previous findings^23^. We subsequently used the estimated decay rates and transformed burst frequencies into hours and found the duration between two bursts of lncRNAs from one allele to be more than twice as long compared to mRNAs (15.9 and 6.9 hours, respectively, median) (**Figures 2G** and **S4G**). Notably, over 30% of lncRNAs were found to burst less than once every 24 hours on each individual allele.

Having determined that lncRNAs show increased cell-to-cell variability compared to expression matched mRNAs (**Figures 1D-G** and **S2A-B**), we next explored if this observation was associated with a systematic pattern of bursting parameters. To this end, we identified the 50 most variable lncRNAs on each allele (ranked CV^2^) (**Figures S4H-I**) and generated thousands of sets of randomly drawn expression-matched mRNAs (similar as in **Figure 1F**). Strikingly, we observed that the lncRNAs with highest CV^2^ had decreased burst frequencies (**Figures 2H** and **S4J**, P < 1e^-4^, permutation test), and increased burst sizes (**Figures 2I** and **S4K**, P=0.015, permutation test) compared to expression matched mRNAs. These data suggest more sporadic expression of lncRNAs (due to lowered burst frequency), although with increased number of RNA molecules produced per burst (due to increased burst size) and link the increased cell-to-cell variability of lncRNAs to a shift in transcriptional bursts.

Many lncRNAs are transcribed in the antisense direction of protein-coding (sense) genes^32^ and we next investigated if such genomic organizations could result in altered transcriptional kinetics. We identified loci with divergent (here referred to the presence of a stable annotated transcript in both sense and antisense direction) mRNA-mRNA pairs (n=1,282), divergent mRNA-lncRNA pairs (n=465) and unidirectional mRNA (n=6,276) transcribed promoters (**Figure S4L**). In line with previous studies^5,33^, we observed increased expression of divergently transcribing promoters (**Figure 2J**), an observation that was consistent for mRNA-mRNA as well as mRNA-lncRNA promoters, compared to unidirectional transcribing promoters (**Figure 2K**). Using the inferred transcriptional kinetics, we next tested whether the increase in expression (**Figure S4M**) was related to an alteration in bursting. We observed a consistent increase in burst frequency for divergently mRNA-mRNA and lncRNA-mRNA transcribing promoters (**Figures 2L** and **S4N**, two-sided Wilcoxon rank-sum test), with no consistent increase in burst size (**Figures 2L and S4N**, two-sided Wilcoxon rank-sum test). The fact that divergent transcription units tend to burst more frequently - although without increased RNA production per burst - likely suggests that two closely situated promoters facilitate recruitment of the required transcriptional machineries. In support of this, previous reports have suggested that divergent promoters harbor increased number of TF motifs compared to nondivergent promoters^5^.

### Precise expression of lncRNAs in transient cellular phenotypes reveals molecular functions

We next explored if scRNA-seq could serve as a tool for functional annotation of lncRNAs. More specifically, we hypothesized that dynamic expression of lncRNAs in transient cellular states carries information to their function, thus applying a revised ‘guilt-by-association’^34^ principle where lncRNA expression is linked to a cellular state. To evaluate this approach, we first sought to identify lncRNAs involved in cell cycle progression. Asynchronously growing mouse fibroblasts (**Figures S1A-C**) were projected into low-dimensional PCA space using the most variable^35^ cell cycle genes^36^ (**Figures S5A-B, Supplementary Table 3**), clustered, and the PCA coordinates were used to fit a principal curve^37^. Cells were aligned to the fit and cell cycle progression was confirmed by comparing the relative expression of a subset of well-established cell cycle genes marking cells in G0, G1, G1S and G2M (**Figure 3A, Figure S5C**). To identify lncRNAs with high expression within specific phases of the cell cycle, we next applied an ANOVA test (FDR < 0.01, Benjamini-Hochberg adjusted) on gene expression over the individual cell cycle phases. The P-values were contrasted against the fold induction in the particular cell cycle phase (**Figure 3B**), revealing a total of 128 lncRNAs with a cell cycle specific expression pattern (fold change > 1.5, adjusted p < 0.01) (**Figure 3B**), of which we selected seven highly ranked candidates for further characterization.

**Figure 3.**
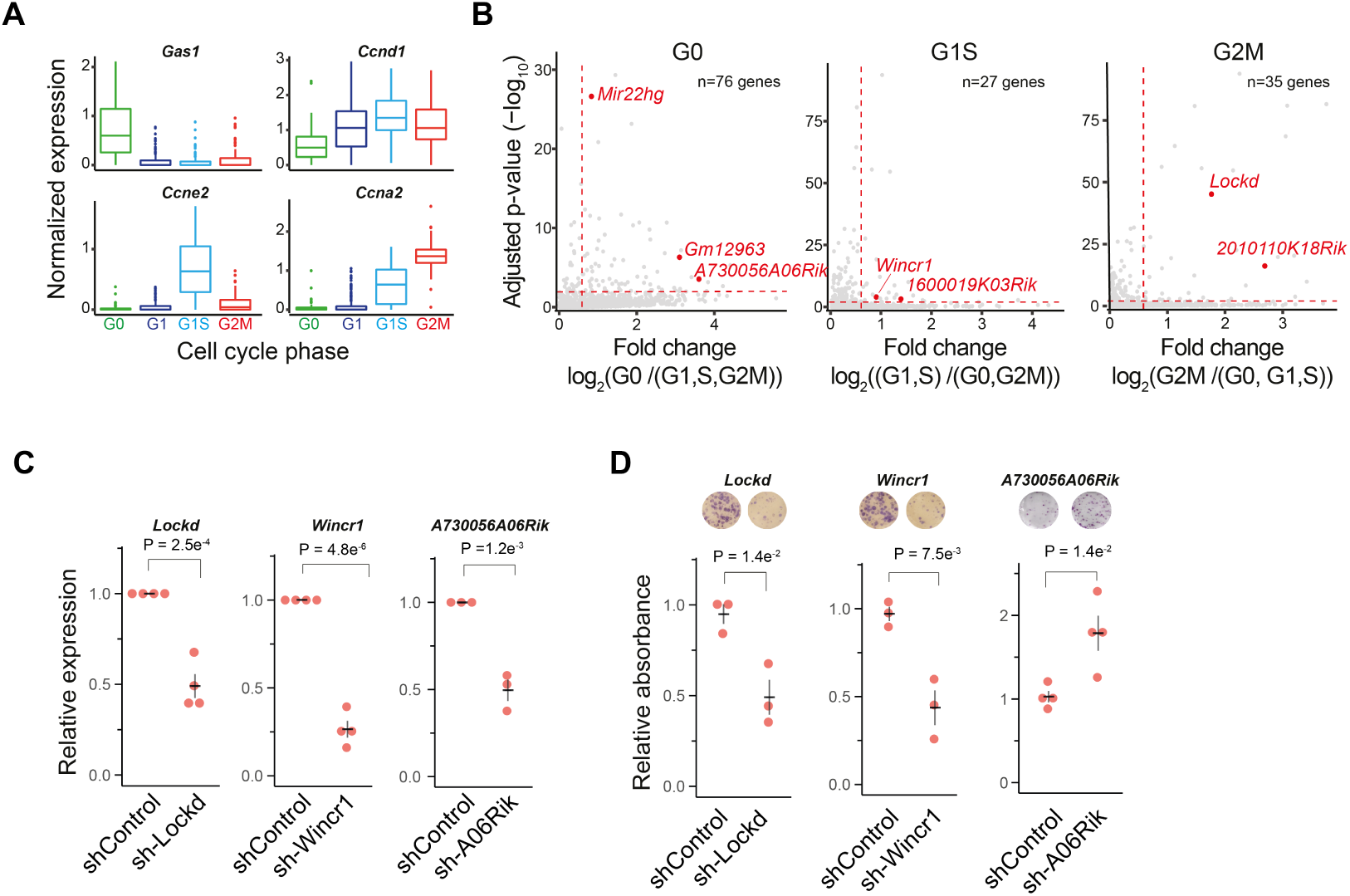
Identification of cell cycle regulated lncRNAs using scRNA-seq. **(A)** Boxplots showing the normalized expression levels (scale-factor and log10 normalized^35^) of cell cycle marker genes in cells classified to cell cycle phase. **(B)** Scatter plots showing lncRNAs with significant expression differences across cell cycle phases (y-axis, Benjamini-Hochberg adjusted ANOVA test) against the fold induction compared to the other cell cycle phases. The top ranked candidates selected for further validation were colored red. **(C)** Relative expression levels of candidate lncRNAs in lentiviral transduced NIH3T3 cells measured by qRTPCR. P-values represent a two-sided student’s t-test. **(D)** Quantification of colony forming cells in sh-control cells and cells with stable shRNA-induced knockdown of lncRNAs, together with representative photos of staining. P-values represent a two-sided student’s t-test.

To functionally evaluate the selected lncRNA candidates’ potential involvement in cell cycle progression, we used the immortalized mouse embryonic fibroblast NIH3T3 cell line. NIH3T3 was found to express similar cell cycle genes^36^ as primary fibroblasts (**Supplementary Table 3**) and also correlate well in expression levels (**Figure S5D**). Next, cell cycle progression of NIH3T3 cells was synchronized by serum starvation (G0/G1), thymidine block (G1S) or nocodazole treatment (G2M), and the synchronization was validated by flow cytometry (**Figure S5E**) and qRTPCR for two cell cycle marker genes (**Figure S5F, Supplementary Table 4**). As expected, we found all seven lncRNAs to have peak expression in the predicted phases of the cell cycle by qRTPCR validation (**Figure S5G**). Having validated cell cycle specific expression of these lncRNA transcripts, we next generated individual lentiviral transduced NIH3T3 cell lines with stable shRNA-induced knockdown for three of the candidates (*Wincr1, Lockd* and *A730056A06Rik*) in order to perform in-depth functional investigation (**Figure 3C**). Although no clear phenotype was observed for these cells under normal growth conditions, striking effects were observed in colony formation assays (**Figure 3D**), which provide a moderate stress on cells. While the knockdown of *A730056A06Rik* (expressed upon serum starvation, **Figure S5G**) resulted in the formation of more colonies, the knockdown of *Wincr1* and *Lockd* (expressed in proliferating cells, **Figure S5G**) reduced the numbers of colonies formed (**Figure 3D**). Together, this showed that lncRNAs selected with our new screening paradigm could be efficiently related to their cellular function.

### Functional investigation of the lncRNA *Lockd*

We were intrigued by a previous report^38^ suggesting that transcription of the *Lockd* gene functions *in cis* by promoting expression of the cell cycle regulator *Cdkn1b* (10kb upstream of the *Lockd* locus, **Figure S6A**) whereas the *Lockd* transcript was found dispensable for this mechanism. Although no function was reported for the *Lockd* transcript in that study^38^, our colony formation experiment showed that reduction of the *Lockd* transcript through shRNA knockdown had a clear functional consequence – by reducing the colony formation capacity of the cells (**Figure 3D**). To complement the stable *Lockd* knockdown, we designed two siRNAs (**Supplementary Table 5**) against the *Lockd* transcript which both achieved good knockdown efficiency with less than 10% *Lockd* expression in the NIH3T3 cell line as well as in primary fibroblasts (**Figure S6B**). In agreement with the NIH3T3-shLockd stable cell line (**Figure 3D**), a consistent decrease in colony forming cells was observed upon siRNA-induced *Lockd* depletion (**Figure 4A**). In line with the previous report^38^, no consistent change in RNA expression was observed for *Cdkn1b* upon knockdown of the *Lockd* transcript in NIH3T3 or primary fibroblast cells (**Figure 4B**). However, the allele-resolved scRNA-seq data suggested co-expression of *Lockd* and *Cdkn1b* (tended to be expressed in the same cells and from the same allele) on both the CAST (Fisher’s exact test, P=0.013) and C57 (Fisher’s exact test, P=0.00032) alleles (**Figure S6C**).

**Figure 4.**
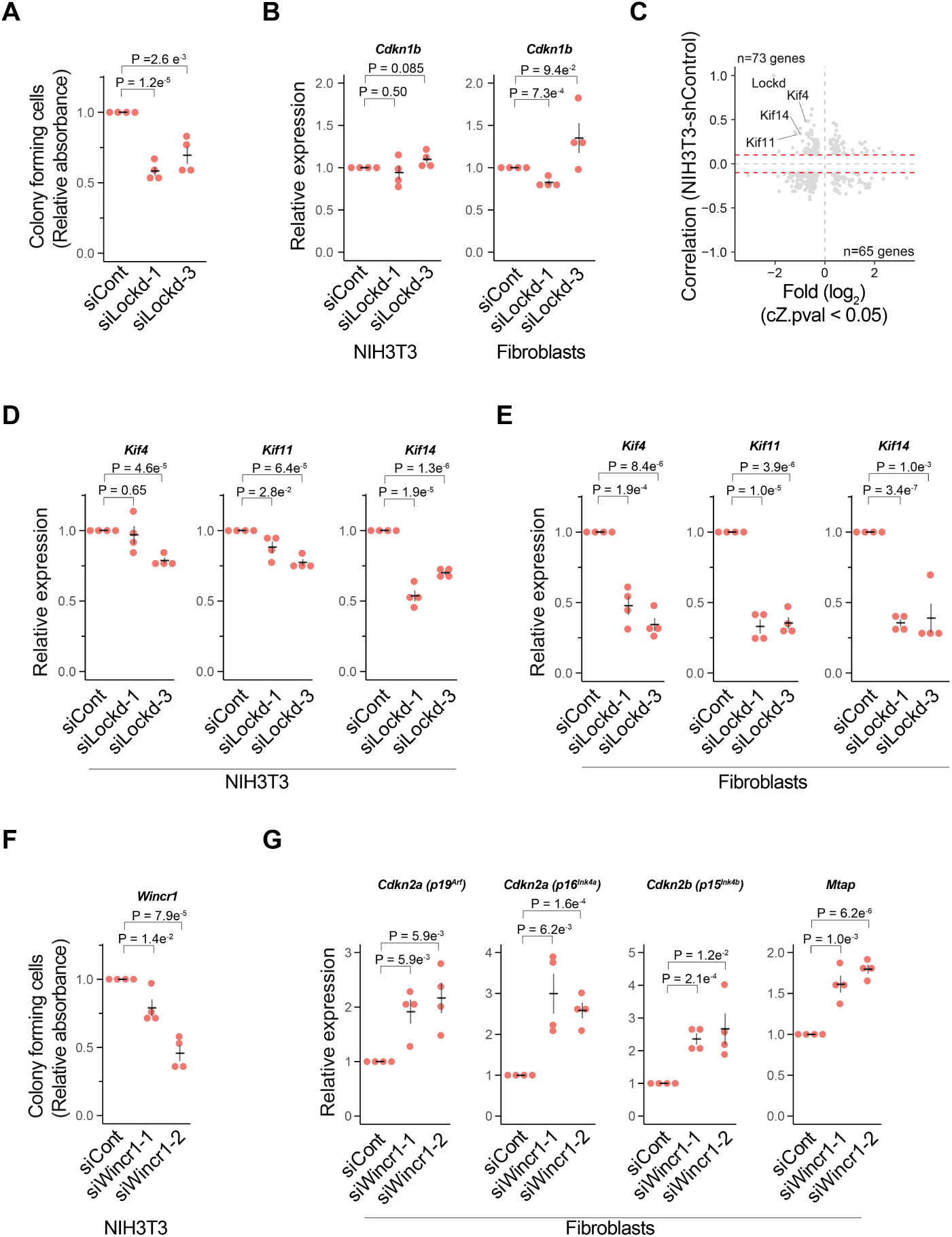
Functional analysis of *Lockd* and *Wincr1* lncRNAs. **A)** Quantification of colony forming cells upon siRNA induced knockdown of *Lockd* in NIH3T3 cells (n=5, p-values represent a two-sided student’s t-test, error bars represent the s.e.m.). **(B)** Relative expression of *Cdkn1b* upon siRNA induced knockdown of *Lockd* in NIH3T3 cells (n=5) and primary fibroblasts (n=4) measured by qRTPCR. P-values represent a two-sided student’s t-test, error bars represent the s.e.m. **(C)** Scatter plot representing magnitudes of fold changes of gene expression (shLockd / shControl, for significant genes with adjusted p-value < 0.05) of stably transduced NIH3T3 cells (x-axis) against gene-gene correlations to *Lockd* in stably shControl-transduced NIH3T3 cells (y-axis). Genes reaching the threshold of Spearman correlations (denoted with red dashed lines) were considered for downstream analysis. **(D)** Relative expression of candidate genes upon siRNA induced knockdown of *Lockd* in NIH3T3 cells measured by qRTPCR (n=4, p-values represent a two-sided student’s t-test, error bars represent the s.e.m.). **(E)** Relative expression of candidate genes upon siRNA induced knockdown of *Lockd* in primary fibroblast cells measured by qRTPCR (n=4, p-values represent a two-sided student’s t-test, error bars represent the s.e.m.). **(F)** Quantification of colony forming cells upon siRNA induced knockdown of *Wincr1* in NIH3T3 cells (n=5, p-values represent a two-sided student’s t-test, error bars represent the s.e.m.). **(G)** Relative expression of candidate *cis*-interacting genes upon siRNA induced knockdown of *Wincr1* in primary fibroblast cells measured by qRTPCR (n=5, p-values represent a two-sided student’s t-test, error bars represent the s.e.m.).

To characterize the molecular function of *Lockd* further, we used scRNA-seq on the shControl (n=147 cells) and shLockd stable cell lines (n=144 cells). We observed that 752 genes had significantly altered expression in the *Lockd* reduced cell line (SCDE^39^, adjusted p<0.05) (**Figure S6D**), likely including both direct effects and indirect effects of impaired cell cycle progression and possible shRNA-induced off-target effects. Next, we calculated Spearman pairwise correlations of expression levels from the shControl treated stable cell line and hypothesized that the most relevant targets for *Lockd* would have a positive correlation with *Lockd* and concurrent reduced expression in the shLockd cells or alternatively, negative correlation with increased expression upon knockdown (**Figure S6E**). This approach strongly reduced the number of candidate genes (752 to 138 genes) and revealed several well-established cell cycle regulators. Particularly, three members of the kinesin superfamily (*Kif4, Kif11* and *Kif14*), a group of proteins known to be involved in mitosis, appeared as main candidates (**Figure 4C**). Notably, a link between these genes and Cdkn1b has been suggested. While Cdkn1b acts as a transcriptional suppressor of *Kif11* by binding to the *Kif11* promoter through a p130-E2F4 dependent mechanism^40^, Kif14 regulates protein levels of Cdkn1b through a proteasome-dependent pathway^41^. Based on these previous findings, we set out to directly confirm the effect on *Kif4, Kif11* and *Kif14* by measuring the expression levels on qRTPCR upon siRNA induced knockdown of *Lockd* in NIH3T3 and primary fibroblast cells. The effect on *Kif11* and *Kif14* was seen in both cell lines while the effect on *Kif4* could only be observed in the primary fibroblasts (**Figures 4D-E**). However, this is consistent with the scRNA-seq data (**Figure 4C**) where *Kif4* was more modestly affected compared to *Kif11* and *Kif14*. In summary, our data suggest *Lockd* to function through *trans* regulatory mechanisms, in addition to its previously reported *cis*-function^38^. While transcription of the *Lockd* gene functions *in cis* to promote transcription of *Cdkn1b*^38^, the *Lockd* transcript appears to function in the same pathway as Cdkn1b and enhances the negative effects on cell cycle progression.

### Functional investigation of the lncRNA *Wincr1*

We next sought to investigate the molecular function of *Wincr1* in greater detail. To this end, two siRNAs were first designed against the *Wincr1* transcript and knockdown confirmed by qRTPCR (**Figure S7A**). In line with the NIH3T3-shWincr1 stable cell line, a decrease in colony forming cells - which scaled with the degree of knockdown in NIH3T3 cells - was observed upon siRNA-induced depletion of *Wincr1* (**Figure 4F**). We next turned to our scRNA-seq data (**Figures S1A-C**) and observed that several genes at this locus, *Cdkn2a (*p16^Ink4a^ and p19^Arf^), *Gm12602* and *Mtap*, presented coordinated expression with *Wincr1* (**Figure S7B**). Intriguingly, the homologous region in human has been reported to regulate the expression of *CDKN2A* (p16^INK4A^) in a mechanism where the microRNA-31 host gene (MIR31HG) recruits chromatin remodeling factors to the promoter of p16^INK4A, 42^. However, Mir31hg has a different genomic structure in mouse while *Wincr1* is absent in human cells (**Figures S7C-D**). We therefore sought to investigate if *Wincr1* had similar functions as human MIR31HG. Indeed, siRNA-induced knockdown of *Wincr1* in primary fibroblasts showed a significant increase in the expression of *Cdkn2a* (p16^Ink4a^ and p19^Arf^) as well as *Cdkn2b* (p15^Ink4b^) and *Mtap* (**Figure 4G**), suggesting *Wincr1* functions *in cis*. Finally, we also noted that *Wincr1* has been proposed to be involved in Wnt/β-catenin signaling and to attenuate cell migration through *trans*-acting mechansisms^43^. Since *Cdkn2a* (p16^Ink4a^ and p19^Arf^)^44^ as well as *Cdkn2b* (p15^Ink4b^)^44^ are inactivated in NIH3T3 cells due to homozygous deletions of their chromosomal regions, our data support that *Wincr1* also maintains other *trans*-acting functions (**Figure 4F**).

### Functional annotation of lncRNAs can be generalized to several phenotypes

We next evaluated if the presented approaches are applicable to other cellular states and set out to investigate whether lncRNAs involved in apoptotic signaling could be identified. Since apoptotic signaling is linked to proliferation, we limited the analysis to cells in the G1 phase (**Figure 3A**) and repeated the low-dimensional projection, now using the most variable genes related to apoptotic signaling (**Figure S8A**). We identified three clusters of cells (**Figure 5A**) and focused specifically on one cluster of cells that expressed genes involved in stress signaling, namely *Gadd45b*^45^ (Growth arrest and DNA-damage-inducible, beta) and the p53 target gene *Cdkn1a* (**Figures 5B and S8B**). Single-cell differential expression (SCDE)^39^ was applied to find lncRNAs with increased expression in this cluster of cells and a total of five highly ranked lncRNAs, towards which siRNAs could be designed, were selected for further validation (**Figure 5C**). To validate these candidates, DNA damage induced apoptosis was triggered in the NIH3T3 cell line by treating cells with the chemotherapeutic and DNA crosslinking reagent Mitomycin C (MMC). DNA damage was validated using qRTPCR by measuring the expression levels of *Cdkn1a* (induced by p53) and *Gadd45b* (**Figure S8C**) and, in line with the prediction, expression of all five candidate lncRNAs were induced upon MMC treatment and scaled with the concentration of MMC (**Figure 5D**). To further investigate the regulatory effects these lncRNA candidates might have on apoptosis, three of the candidates were suppressed by two siRNAs each (**Figure S8D**). The levels of apoptosis in lncRNA suppressed NIH3T3 cells was measured by AnnexinV on flow cytometry after treatment with MMC (**Figure 5E**). Notably, apoptosis was repeatedly induced when exposed to MMC, thus suggesting that knockdown of these lncRNAs sensitize cells to undergo apoptosis. In summary, our data support that the effect of lncRNAs on more subtle cellular states, such as pro-apoptotic signaling, are also captured by scRNA-seq and serves as a profound tool to link lncRNAs to function.

**Figure 5.**
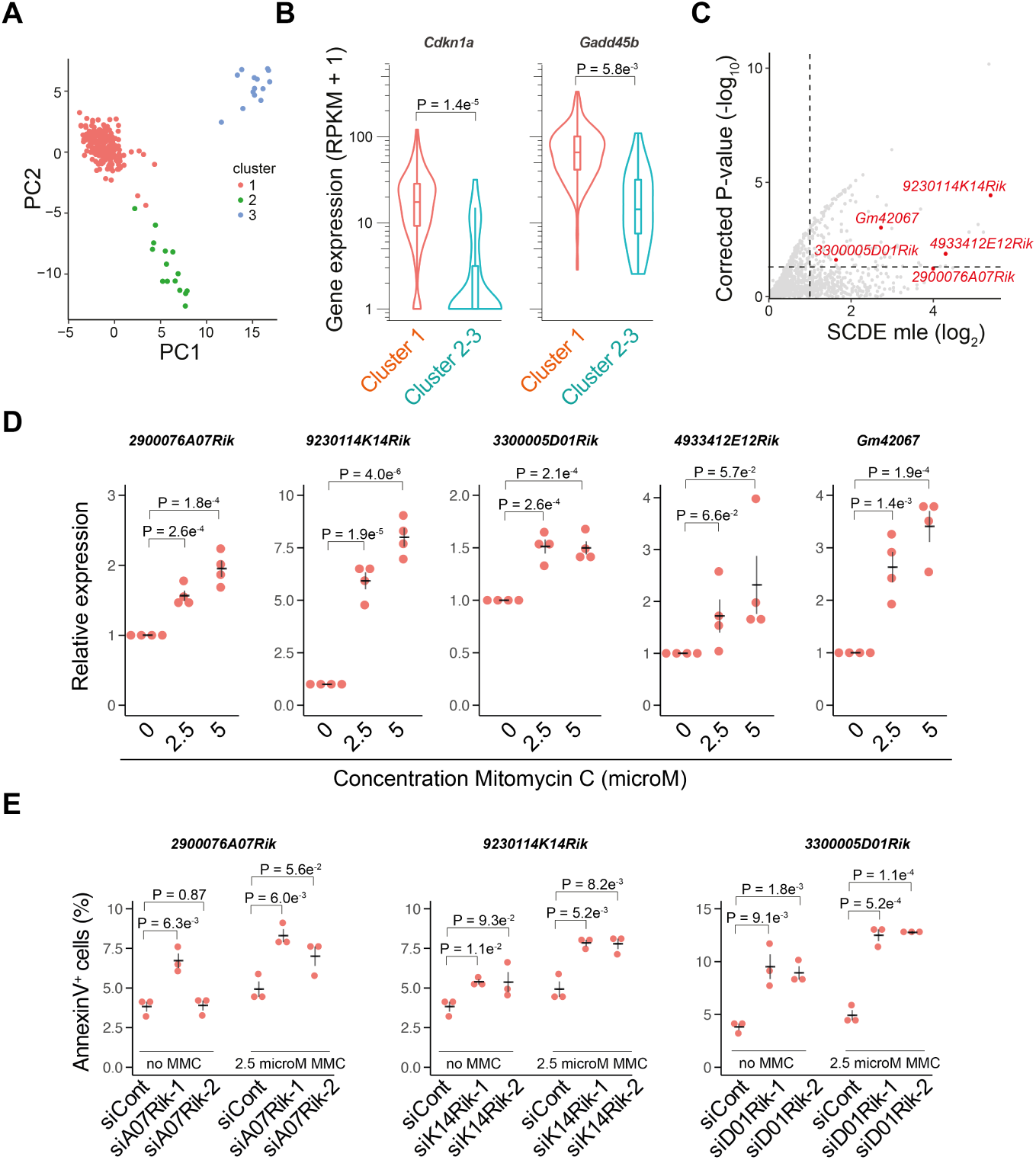
Identification of lncRNAs involved in apoptosis by single-cell profiling. **(A)** Low dimensional PCA projection of cells, based on the most variable genes annotated as apoptosis related. Cells are colored according to clusters (identified by the pam function of the R package ‘cluster’). **(B)** Violin plots showing the expression levels of two marker genes for DNA damage (P-values represent a two-sided Wilcoxon rank-sum test). **(C)** Scatter plot showing fold change magnitudes (x-axis) and significance levels (y-axis) for cluster 1 against clusters 2-3 identified in (A) analyzed by SCDE^39^. lncRNAs selected for validation are marked in red. **(D)** Relative expression measured by qRTPCR of candidate lncRNAs in NIH3T3 cells treated with MMC (n=4, p-values represents a two-sided student’s t-test, error bars represent the s.e.m.). **(E)** Quantification of apoptosis using AnnexinV for siRNA targeted NIH3T3 cells treated with MMC (n=3, p-values represents a two-sided student’s t-test, error bars represent the s.e.m.).

### Allele-sensitive expression reveals lncRNA-mRNA interacting gene pairs

Previous bulk RNA-seq studies identified pervasive allelic imbalance of gene expression across heterozygous F1 hybrid mice^46^. We next asked if allelic imbalance of lncRNAs could reveal information about *cis* regulatory mechanisms and gene-gene interactions (**Figure 6A**). To improve the power to detect gene-gene interactions, we profiled additional 218 mouse adult tail fibroblasts of the reciprocal cross (CASTxC57), resulting in a comprehensive Smart-seq2 dataset of 751 cells (post quality control, ∼4 Million mapped reads per cell, median) containing multiple independent mouse explant cultures (**Figures S9A-C and S1A-C**). We calculated allelic distributions by counting allele sensitive read counts over both genomes and quantified the allelic imbalance as (CAST_allelicCounts_ /(CAST_allelicCounts_ + C57_allelicCounts_) - 0.5) where a positive score reflects increased RNA expression towards the CAST genome. Consistent with previous bulk RNA-seq studies^46^, we confirmed ∼75% of mouse genes (8,981 of 11,377) to have RNA expression levels dependent on the genetic background (**Figure S9D**, binomial test, Benjamini-Hochberg adjusted p<0.01) and found lncRNAs to have greater allelic imbalance than mRNAs (**Figure S9E**) across a wide range of expression levels (**Figure S9F**). To identify *cis*-functioning lncRNAs, we first retrieved all lncRNA-mRNA gene pairs within +/- 500 kb of the lncRNA transcript start site (TSS) (5,824 pairs in total, **Figure 6B**) and calculated a score for allelic imbalance for each lncRNA-mRNA gene pair (see Methods). Next, a permutation test was applied, where each lncRNA was moved to 1,000 randomly selected gene locations and the score for *in silico* sampled gene pairs recomputed (+/- 500 kb of the lncRNA TSS, 6.75e^6^ random gene pairs in total, **Figure 6C**). In total, 90 significant lncRNA-mRNA interactions were identified (p<0.05, permutation test, **Supplementary Table 6**) and significant gene pairs found to be enriched at closer distance (within 25 kb, **Figures 6D-E**). We then sorted the significant interactions using the score of allelic imbalances where allelic imbalance towards the same allele was assigned a positive score, and allelic imbalance on opposite alleles a negative score (**Figure 6F**). Four highly ranked lncRNA-mRNA interactions, all accessible to siRNA depletion and with diverse genomic organization (**Figure S9G**), were examined more deeply. Upon evaluation of allele-informative reads for all genes within 500 kb of the lncRNA TSSs, few other genes (only one) were found to have any pronounced allelic imbalance (**Figure S9H** and **Supplementary Table 6**), therefore suggesting these lncRNAs predominantly interact with only individual mRNAs at their loci.

**Figure 6.**
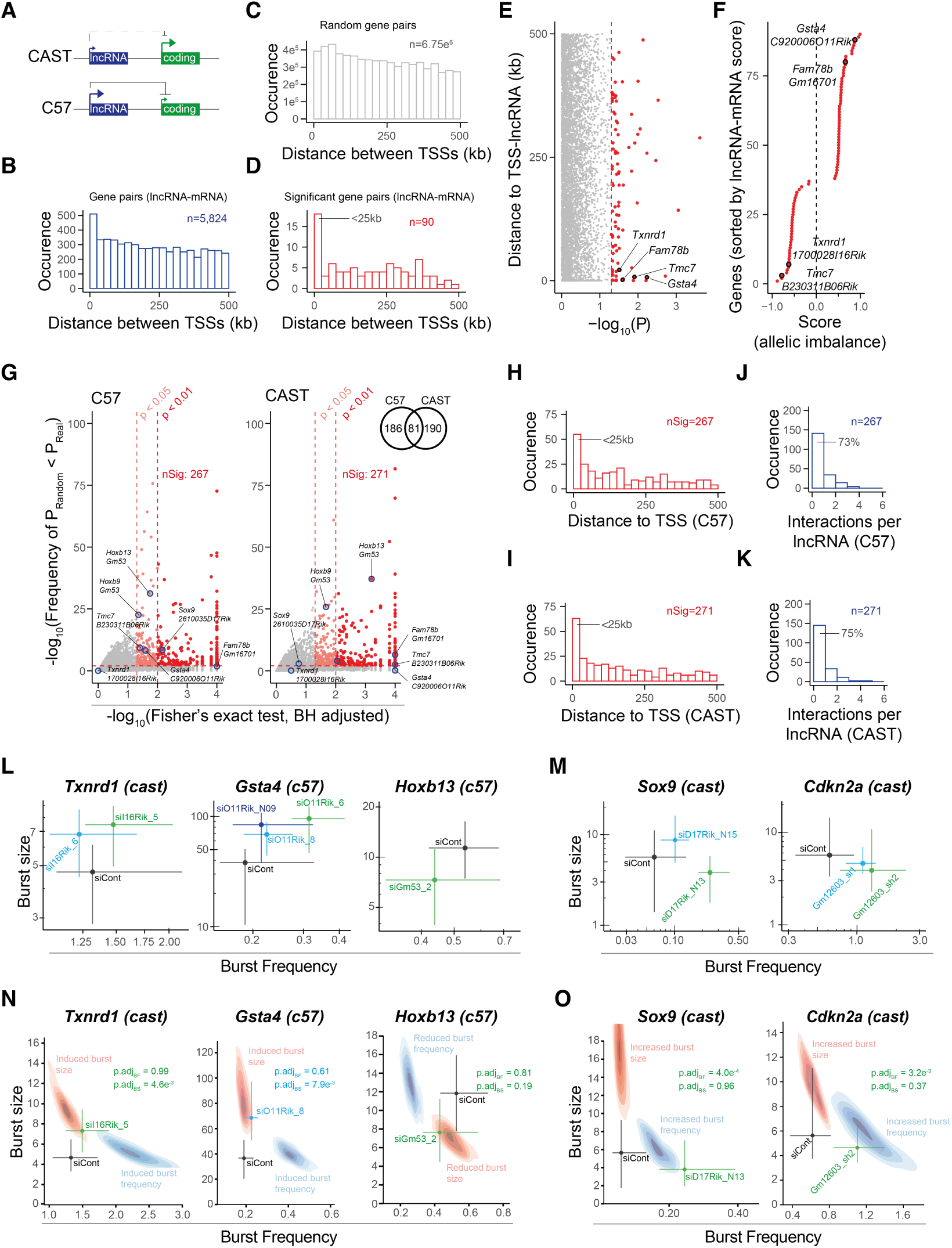
Identification of *cis* functioning lncRNAs using allele-resolved expression. **(A)** Schematic illustration of hypothetical *cis*-effects between lncRNAs and proximal mRNAs that could manifest as coordinated allelic expression dynamics across cells. **(B)** Histogram showing lncRNA-mRNA gene pairs within +/- 500 kb of lncRNA TSSs. **(C)** Histogram showing permutated lncRNA-mRNA gene pairs (where each lncRNA was moved to 1,000 randomly selected gene locations). **(D)** Histogram showing significant lncRNA-mRNA gene pairs (p < 0.05, permutation test). **(E)** Scatter plot representing p-values (permutation test) against the distance between lncRNA-mRNA pairs of genes. Significant gene pairs are colored in red, dashed red line represents p=0.05. **(F)** Scatter plot representing ranking of significant lncRNA-mRNA pairs of genes. A positive value (x-axis) represents allelic bias towards the same allele while a negative value represents bias on opposite alleles. Highlighted lncRNA-mRNA pairs of genes are considered for downstream validation. **(G)** Scatter plot representing p-values (permutation test) against the distance between lncRNA-mRNA pairs of genes. Significant gene pairs are colored in red. Dashed red line represents p=0.01, dashed light red line represents p=0.05. Venn diagram represents significant lncRNA-mRNA pairs of genes for the C57 and CAST genomes. **(H-I)** Histogram showing significant lncRNA-mRNA pairs of genes for the (H) C57 and (I) CAST genome. **(J-K)** Histogram representing the number of significant interactions for individual lncRNAs for the (J) C57 and (K) CAST genome. **(L)** Scatter plots representing burst parameters with 95% confidence intervals for *Txnrd1, Gsta4* and *Hoxb13* upon siRNA induced knockdown of lncRNAs. **(M)** Scatter plots representing burst parameters with 95% confidence intervals for *Sox9* and *Cdkn2a* upon siRNA induced knockdown of lncRNAs. **(N)** Scatter plots representing burst parameters with 95% confidence intervals for *Txnrd1, Gsta4* and *Hoxb13* upon siRNA induced knockdown of lncRNAs for one representative siRNA from Figure 6L. Simulated cases when expression is modulated by burst frequency or size are shown in blue and red, respectively. P-values represent a profile likelihood test. **(O)** Scatter plots representing burst parameters with 95% confidence intervals for *Sox9* and *Cdkn2a* upon siRNA induced knockdown of lncRNAs for one representative siRNA from Figure 6M. Simulated cases when expression is modulated by burst frequency or size are shown in blue and red, respectively. P-values represent a profile likelihood test.

To fully take advantage of the allelic resolution, we next assessed allele-specific expression patterns on the single-cell level for the same set of lncRNA-mRNAs gene pairs as above (5,824 gene pairs +/- 500 kb of the lncRNA TSS, **Figure 6B**). To identify the most significant candidates, a Fisher’s exact test was first applied on each gene pair (P_Real_, Benjamini-Hochberg adjusted) and also for *in silico* sampled gene pairs by moving each lncRNA to 1,000 randomly selected gene locations (P_Random_, Benjamini-Hochberg adjusted, similar as in **Figure 6C-E**). For downstream analysis, we primarily considered genes with P_Real_ < 0.01 where P_Random_ < P_Real_ occurred in less than 1% of the permutated gene interactions (**Supplementary Tables 7** and **8**). These criteria identified 457 lncRNA-mRNA gene pairs with significant coordinated expression on at least one of the alleles (**Figures 6G**). Significant interactions were again found to be enriched at closer distance (<25 kb, **Figures 6H-I)** and most lncRNAs observed to interact with only one mRNA (**Figures J-K**). Encouraged to see that several of the candidates overlapped between the two approaches (**Figures 6E and 6G**), we next sought to functionally dissect a subset of significant interactions. To this end, six lncRNA-mRNA gene pairs were selected for further validation, including two that were identified by both approaches (*B230311B06Rik:Tmc7* and *Gm16701:Fam78b*), two using allelic imbalance (*1700028I16Rik:Txnrd1* and *C920006O11Rik:Gsta4*) and two that were identified by our single-cell resolution (*2610035D17Rik:Sox9* and *Gm53:Hoxb13*). We also noted that the lncRNA *Gm53* showed a second significant interaction with *Hoxb9* (in addition to *Hoxb13*) at a slightly lower significance threshold (P_Real_ < 0.05) (**Figure 6G**). To evaluate these molecular interactions, we designed at least two siRNAs against each lncRNA and measured the effects with qRTPCR. All candidate gene pairs were confirmed to show the expected target mRNA expression change (**Figures S10A-F**), and we also validated an increase in unspliced RNA levels for *Gsta4* and *Txnrd1*, thus indicating that the regulatory interplay acts on the transcriptional level (**Figures S10A-B)**.

While it has been clearly established that many lncRNAs effect transcription of target mRNAs, it is not known if lncRNAs act by altering their burst frequencies or sizes. To address this question, we turned to our validated lncRNA-mRNA interactions (*1700028I16Rik:Txnrd1, C920006O11Rik:Gsta4, Gm16701:Fam78b, B230311B06Rik:Tmc7, 2610035D17Rik:Sox9, Gm53:Hoxb9, Gm53:Hoxb13* **(Figure S10)** and *Wincr1:Cdkn2a* (**Figure 4G**)) which all had mRNA targets expressed in a part of the parameter space with good precision (narrow confidence intervals, see Methods) for burst inference (**Figures S11A-B**). To obtain burst parameters across lncRNAs perturbations, we profiled individual adult tail fibroblasts with Smart-seq3^22^ upon siRNA induced knockdown and generated a comprehensive data set with at least 200 cells (post quality control) for each siRNA knockdown (**Figures S11C-F**). We first compared fold changes of the Smart-seq3 measurements (**Figures S11G-L**) with those of qRTPCR (**Figures S10A-F**) and found good agreement. Noteworthy, knockdown of lncRNA-Gm53 using siGm53_3 was found less efficient than siGm53_2 on both qRTPCR (**Figures S10D)** and scRNA-seq measurements **(Figure S11M**) and the induction on *Txnrd1* was less robust for siI16Rik_6 compared to siI16Rik_5 (**Figure S11H**). We next inferred burst parameters for *Txnrd1, Gsta4, Sox9, Cdkn2a* and *Hoxb13* while *Tmc7* and *Fam78b* did not reach sufficient UMI counts and SNP coverage for burst inference. The inference showed a consistent effect on burst size for *Txnrd1, Gsta4* and *Hoxb13* (**Figure 6L**), whereas *Sox9* and *Cdkn2a* showed an increase in burst frequency (**Figure 6M**). Using simulations for one representative siRNA for each lncRNA, we demonstrated that the observed effects were in the regions of parameter space expected for an exclusive effect on either burst size (**Figure 6N**) or burst frequency (**Figure 6O**). Taken together, these observations suggest that lncRNAs can regulate both burst frequencies and burst sizes, possibly through independent molecular processes.

## Discussion

The genomic sequences that harbor transcriptional activity and give rise to lncRNAs have been comprehensively mapped^4^. Yet it has remained hard to determine lncRNA functions, and it has been proposed that their low expression^4–9^ is one of the main challenges for functional characterization. Here, we detected a typical lncRNA in only 3% of cells (median, **Figure 1C**) and leveraged our single-cell approach to map lncRNA functions by systematic characterization of these rare cells. More precisely, we here sought to map lncRNA burst characteristics and functions using scRNA-seq followed by functional validation experiments.

It has long been accepted that lncRNAs are expressed at lower levels than mRNAs but the underlying molecular causes have remained unclear^4^. Using allele-resolved scRNA-seq, we contrasted the transcriptional burst kinetics for several hundreds of lncRNA genes with those of mRNAs. We discovered that low expression of lncRNAs is mainly governed by lowered burst frequencies (**Figures 2D** and **S4A**) and found the duration between two transcriptional bursts of lncRNAs to be approximately twice as long compared to mRNAs (**Figures 2G** and **S4G**). Notably, over 30% of lncRNAs were estimated to burst less than once every 24 hours from each allele, suggesting that many lncRNA alleles may remain inactive throughout an entire cell cycle. While the lowered burst frequency of lncRNAs (four-fold decrease, **Figures 2D** and **S4A**) likely represent a decrease in enhancer-mediated transcriptional initiation, the more modest effect on burst size (two-fold decrease, **Figures 2E and S4B**) could relate to differences in promoter features.

Interestingly, loci with two genes divergently transcribed from the same promoter regions (lncRNA-mRNA as well as mRNA-mRNA gene pairs) had higher burst frequencies than genes separated by larger distances (**Figures 2L and S4N**). The increased burst frequency might stem from promiscuous interactions between regulators at the closely located promoters so that both promoters can more easily load the core transcriptional complexes. It is also possible that the presence of more TF binding sites^5^ at divergent promoters could result in increased burst frequency.

We also revisited the question whether lncRNAs have increased cell-to-cell variability in expression compared to mRNAs of similar expression. Although lncRNA expression patterns are heterogeneous (**Figures 1D-E**), we observed in both mouse and human model systems that lncRNAs had generally higher cell-to-cell variability (quantified as CV^2^) compared to mRNAs of the same average expression level (**Figures 1E and S2A-B**). It is possible that the larger number of lncRNAs monitored with the scRNA-seq data presented in this study, explains why earlier reports with lower gene counts did not identify any increased variability of lncRNAs^15^. In line with this argument, subsampling lncRNA loci from our scRNA-seq data showed declining and eventually complete loss of power to detect the variability difference between lncRNAs and mRNAs (**Figure 1H**). We compared burst kinetics of lncRNAs with expression matched mRNAs and found that the increase in cell-to-cell variability associated with a shift in burst parameters where some lncRNAs burst less frequently although with higher burst sizes (**Figures 2H-I** and **S4J-K**). Thus, our observations support previous models suggesting lncRNAs to be expressed in few cells (more sporadic expression in cells over time), although at slightly higher levels^14^, for at least a subset of lncRNAs.

Analysis of scRNA-seq data also allowed us to identify *cis-* and *trans-* functions of lncRNAs. By assigning expression of lncRNAs to transient cellular states, such as different cell cycle stages and pro-apoptotic signaling, we applied a revised ‘guilt-by-association’^34^ principle where functions of lncRNAs was linked to a particular cellular condition. We revealed that this approach can generate hypotheses of lncRNA-functions without the need for perturbation experiments. Notably, knockdown of several candidate lncRNAs did not have a clear phenotype (data not shown) until exposed to a relevant stress stimulus (**Figure 3D**) and are therefore likely prone to be missed in large genome-wide perturbation studies carried out in steady-state growth conditions. Finally, we also found that overlapping of gene-gene expression correlations from scRNA-seq data with differential expression upon knockdown of lncRNAs, is a highly useful strategy for decoding lncRNA functions (**Figures 4C** and **S6E**). We believe this approach reduced the background significantly and exposed the most relevant targets for our studies on *Lockd* (**Figure 4C**).

We finally explored how lncRNAs modulate burst kinetics of nearby protein-coding genes. Although it has been clearly recognized that many lncRNAs function as transcriptional regulators^10,11,47^, a major gap in our understanding has been how this influence transcriptional bursts. Regulation of burst dynamics is generally poorly understood, and we here provide evidence that lncRNAs can modulate both burst sizes and burst frequencies (**Figures 6L-O**). Clearly, more lncRNA-mRNA interactions need to be characterized in greater detail to fully capture and generalize molecular mechanisms. Yet, our observations imply that lncRNAs modulate enhancer-controlled initiation frequencies of transcription (by modulating burst frequency **Figure 6M**) or the numbers of RNA polymerase II complexes that gets loaded during an active burst (by modulating burst size, **Figure 6L**), possibly representing distinct molecular pathways of lncRNA-mediated regulation. Unfortunately, several of the evaluated lncRNA-mRNA gene pairs (two out of seven) did not provide enough precision to determine a clear effect on burst parameters. Notably, precision of the inferred burst parameters are gene specific (**Figures S11A-B**) and is affected by gene expression levels, SNP coverage, the number of cells sequenced and the sequencing depth of the experiments. The development of more sensitive scRNA-seq protocols, lowered cost for sequencing and a general increased throughput of cells, should improve the precision in burst inference and allow for analysis at larger scale in future studies.

In summary, our study demonstrates that allele-sensitive scRNA-seq reveals functional signatures of lncRNAs. We here introduce computational and experimental strategies to link lncRNAs functions acting in *trans* and *cis*. We anticipate that the strategies presented here, together with the increasing amount of scRNA-seq data becoming available from different tissues, diseases and species, will advance and facilitate functional studies of lncRNAs. It is also tempting to speculate that scRNA-seq could be particular effective to identify lncRNA functions since the RNA itself is the molecular effector, in contrast to mRNAs, where translation occurs before reaching the functional entity.

## Acknowledgements

This work was supported by grants to R.S. from the Swedish Research Council (2017-01062), the Knut and Alice Wallenberg Foundation (2017.0110), the Göran Gustafsson Foundation, the National Institute of Health (R01MH109556), the Bert L. and N. Kuggie Vallee Foundation and the Ludwig Cancer Research, Stockholm branch. The NIH3T3 cell line was a kind gift of Marianne Farnebo (Karolinska Institutet).

## Author contributions

P.J. designed experiments, sequenced transcriptomes of cells, performed computational analysis, prepared figures and wrote the manuscript. C.Z. sequenced transcriptomes of cells and provided support on computational analysis. L.H. performed experiments. G.J.H. cultured primary fibroblast cells. M.H.J. provided support on Smart-seq2 and Smart-seq3 protocols. B.R. provided support on computational analysis. R.S. designed experiments, supervised the work and wrote the manuscript.

## Data availability

Single-cell RNA-sequencing data generated in this study is being deposited at Array Express at European Bioinformatics Institute: Smart-seq2 data on fibroblasts (ID pending) and stable lentiviral transduced shRNA NIH3T3 cells (ID pending), Smart-seq3 data on fibroblasts (ID pending) and lncRNA knockdown experiments (ID pending). Bulk RNA-sequencing data on actinomycin D treated fibroblasts is being deposited at Array Express at European Bioinformatics Institute (ID pending).

## Code availability

Code for processing scRNA-seq data with zUMIs, kinetic inference, and plotting scripts in R is available upon request.

## Material and methods

### Cell culture

Mouse primary fibroblasts were derived from adult CAST/EiJ x C57BL/6J or C57BL/6J x CAST/EiJ mice (approval by the Swedish Board of Agriculture, Jordbruksverket: N343/12) by skinning, mincing and culturing tail explants in DMEM high glucose (Thermo Fisher Scientific), 10% ES cell FBS (Thermo Fisher Scientific), 1% Penicillin/Streptomycin (Thermo Fisher Scientific), 1% Non-essential amino acids (Thermo Fisher Scientific), 1% Sodium-Pyruvate (Thermo Fisher Scientific), 0.1mM bMercaptoethanol (Sigma) in culture dishes coated with 0.2% gelatin (Sigma). NIH3T3 cells were maintained in DMEM supplemented with 10% FBS (Thermo Fisher Scientific) and 1% Penicillin/Streptomycin (Thermo Fisher Scientific). HEK293FT cells for production of viral particles were maintained in DMEM supplemented with 10% FBS (Thermo Fisher Scientific), 1% Penicillin/Streptomycin (Thermo Fisher Scientific), 1% Non-essential amino acids (Thermo Fisher Scientific), 1% Sodium-Pyruvate (Thermo Fisher Scientific), 0.1mM NEAA (Thermo Fisher Scientific) and 2mM L-Glut (Thermo Fisher Scientific). All cells were cultured in 5% CO_2_ at 37°C.

### Generation of Smart-seq2 libraries

Smart-seq2 libraries were prepared as described earlier^48^ using the following parameters; 1) 20 cycles of PCR for pre-amplification, 2) a ratio of 0.8:1 for bead:sample purification of pre-amplified cDNA (using in-house-produced 22% PEG beads), 3) tagmentation of ∼1 ng of bead purified cDNA using in-house-generated Tn5^49^, 4) 10 cycles of PCR for library amplification of the tagmented samples using Nextera XT Index primers and 5) a ratio of 1:1 for bead purification of DNA sequencing libraries (using in-house-produced 22% PEG beads). Sequencing was carried out on an Illumina HiSeq 2000 generating 43 bp single-end reads.

### Generation of Smart-seq3 libraries

Smart-seq3 libraries were generated according to previously published protocol^22^. Briefly, primary mouse fibroblasts were obtained from tail explants of CAST/EiJ × C57/Bl6J mice (>10 weeks old) and passaged for at least ten days. For knockdown of lncRNAs, cells were seeded at 60,000 cells / well in 6 well plates, transfected at a final concentration of 10nM siRNAs (Lipofectamine RNAiMAX Transfection Reagent), and prepared for sorting 72 hours after transfections. For sorting, cells were stained with propidium iodide before being distributed (using a BD FACSMelody 100 μM nozzle, BD Biosciences) into 384 well plates containing 3 μl of Smart-seq3 lysis buffer (5% PEG (Sigma), 0.10% Triton X-100 (Sigma), 0.5 U μL^-1^ of recombinant RNase inhibitor, (Takara), 0.5 μM Smart-seq3 oligo-dT primer (5’-biotin-ACGAGCATCAGCAGCATACGA T30VN-3’; IDT), 0.5mM dNTP (Thermo Scientific)), spun down and stored at −80 °C immediately after sorting. From this point, standard protocol for Smart-seq3 was applied, using the following parameters; 1) 20 cycles of PCR for pre-amplification of cDNA, 2) a ratio of 0.6:1 for bead:sample purification of pre-amplified cDNA (using homemade 22% PEG beads), 3) tagmentation of 150 ng bead purified cDNA using 0.1 μL of ATM and 4) 12 cycles of PCR for library amplification of the tagmented samples using custom-designed Nextera index primers containing 10-bp indexes. Samples were finally pooled, bead purified at a ratio of 0.7:1 (using homemade 22% PEG beads) and prepared for sequencing on a DNBSEQ-G400RS (MGI) generating 100 bp paired-end reads.

### Processing of RNA-seq data

A subset of primary fibroblasts analyzed in this study (sequenced by Smart-seq2) are part of previously published studies and reanalyzed for consistency^20,26^ present in NCBI SRA (SRP066963). Here, the zUMIs v2.7.1b pipeline^50^ was used for alignment (mouse mm10 assembly), gene quantification (Ensembl gene annotations, GRCm38.91) and allelic calling for primary fibroblasts data. To pass quality control, cells were required to have **1)** >= 500,000 reads, **2)** 4,000 genes expressed at >= 5 read counts **3)** distribution of allelic counts within 0.40 < allelic SNPs < 0.60 on autosomes (imprinted genes and all genes on the X-chromosome excluded, **Supplementary Table 1**) and **4)** no more than 20% of allelic counts mapped to the imprinted X-chromosome (escapee genes excluded, **Supplementary Table 2**). Genes with at least 5 read counts in 2 cells were assigned as expressed and kept for downstream analysis.

Smart-seq3 libraries of HEK293 cells had previously been generated by *Hagemann-Jensen et al*^22^ and deposited in ArrayExpress (E-MTAB-8735). The zUMIs v2.7.0a pipeline^50^ was used for alignment (human hg38 assembly) and quantification of gene expression (Ensembl gene annotations, GRCh38.95). Cells were required to have; **1)** >= 500,000 read counts mapped to exons / cell **2)** >= 500,000 UMI counts / cell and **3)** > = 7,500 genes / cell (>= 1 read count). Genes with at least 1 read count in 3 cells were considered for downstream analysis

Smart-seq2 libraries of mES cells had previously been generated by *Ziegenhain et al*^25^ and is available at Gene expression omnibus (GSE75790). The zUMIs v2.7.2a pipeline was used for alignment (mouse mm10 assembly) and quantification of gene expression (Ensembl gene annotations, GRCm38.91). Cells were required to have; **1)** >= 400,000 read counts mapped to exons / cell and **2)** > = 8,000 genes / cell (>= 5 read counts). Genes with at least 5 read counts in 2 cells were considered for downstream analysis

For Smart-seq3 libraries of primary fibroblasts treated with siRNAs, the zUMIs v2.9.4b pipeline^50^ was used for alignment (human hg38 assembly) and quantification of gene expression (Ensembl gene annotations, GRCh38.95). Cells were required to have; **1)** >= 100,000 read counts mapped to exons / cell and **2)** >= 50,000 unique UMI counts and **3)** >= 5,000 genes (>= 1 UMI counts). Genes with at least 1 UMI read count in 3 cells were considered for downstream analysis.

### Annotation of lncRNAs

The Ensembl BioMart annotations (GRCm38.p6) were used for identification of different subsets of lncRNAs. The BioMart annotations were first filtered for genes passing the quality control thresholds (see above) and lncRNAs (according to BioMart annotations) were categorized as; 1) *Divergent* (no gene-gene overlap and TSSs not separated by more than 500 bp), 2) *Convergent* (gene-gene overlap and TSSs not separated by more than 2 kb, 3) *Intergenic* (no gene-gene overlap and at least 4 kb from any other expressed gene), 4) *Independent transcriptional units* (TSSs separated with at least 4,000 bp from any other expressed gene). The threshold of 4 kb was established by manual inspection of **Figure S1D** where the mean expression had been measured (median of sliding window, size = 51) against the distance between the two most closely located TSSs (only genes passing the quality control was considered for these analysis).

### Permutation test for CV^2^

For the analysis of cell-to-cell variability, only genes meeting the following criteria were considered; **1)** Not imprinted, **2)** Not encoded on the X-chromosome and **3)** being classified as separated transcriptional units (**Figure S1E**).

#### CV^2^

For each lncRNA meeting the criteria, 10 *independently transcribed* protein-coding genes having the most similar mean expression {min(|mean(RPKM_lncRNA_) – mean(RPKM_mRNA_|)} were selected. The matching allowed for the same protein-coding gene to be selected multiple times (sample replacement). For the permutation test (n=10,000), one expression-matched protein-coding gene was randomly sampled for each lncRNA and the expected CV^2^ (median) was calculated for each permutation. The P-value represents the frequency of; median(CV^2^_sampled_) > median(CV^2^_lncRNA_).

For estimating the number of lncRNAs needed for detection of median(CV^2^_lncRNA_) > median(CV^2^_mRNA_), the permutation test was repeat 100 times for each size of subsampling (between 10 and 200 lncRNAs) of the frequency where 50% and 95% of the permutations reached median(CV^2^_lncRNA_) > median(CV^2^_sampled_) was assessed.

### Transcriptional inference of bursting parameters

UMI expression values from Smart-seq3 libraries^51^ were used for transcriptional inference^20^. Briefly, allele-sensitive read counts were used to assign molecule counts (UMIs) to the CAST or C57 genome. Cells having UMIs although lacking allelic read counts for individual genes were assigned as missing values for the inference while cells lacking UMIs as well as allelic information were considered as ‘true’ zeros and included in the analysis. For quality control, genes were required to meet the following criteria; **1)** 1 UMI counts in at least 5 cells, **2)** burst size within 0.2 < size < 50, **3)** burst frequency 0.01 < frequency < 30, **4)** UMI mean expression 0.01 < UMI_mean_ < 100 and **5)** width of confidence intervals (CI_High_ / CI_Low_) below 10^1.5^ (for burst size and frequency). Only non-imprinted autosomal genes, identified as independent transcriptional units, were considered for downstream analysis.

### Identification of cell cycle stage of individual primary fibroblasts

The most variable genes were identified using the R package ‘Seurat’^35^. Genes were first filtered for being expressed in at least 5 cells at 5 read counts. Counts were next normalized using ‘LogNormalize’ (setting scale.factor = 10,000) and the most variable genes were identified using the ‘vst’ method of ‘FindVariableFeatures’. We next extracted the cell cycle related genes reported by Whitfield and colleagues^36^ (**Supplementary table 3**), and used the top 50 ranked genes with the highest variability for PCA. The cell cycle phase of individual cells was identified using the first three principal components as input for the R package ‘princurve’ and the lambda factor used to align individual cells along the cell cycle progression. Expression of individual genes were illustrated using a rolling mean of 15 cells (using R package ‘zoo’). The assignment of cells to cell cycle phase was performed based on the expression levels of known cell cycle regulators (*Gas1, Ccne2, Ccnb1* and *Ccnd1*) using rolling mean of Seurat normalized counts.

### Differential expression analysis of lncRNAs in the cell cycle

Differential expression analysis between cell cycle phases (G0, G1, G1S and G2M) was performed using a one-way ANOVA test (Benjamini-Hochberg adjusted, p<0.01) with normalized counts (lognormalized, Seurat).

### Correlation of cell cycle genes

Genes were first filtered for being expressed in at least 2 cells at 5 read counts for primary tail fibroblasts and GFP+ shControl transduced NIH3T3 cells. The R package ‘Seurat’ was used to log-normalize the read counts and the normalized counts were used to calculate Spearman correlation of cell cycle genes^36^. For each pairwise comparison, cells lacking expression of both genes were excluded from the analysis.

### Cell cycle synchronization

NIH3T3 cells (∼50% confluent) were washed twice in PBS and treated either with 0.1% FBS, 2mM Thymidine or 800nM Nocodazole for 16-24 hours.

### Cell cycle analysis

Cells (including supernatants) were harvested using TrypLE Express (Thermo Fisher Scientific), washed once in PBS, resuspended and fixed in 70% EtOH and stored at -20°C until analysis by flow cytometry. Prior to analysis, cells were washed once in PBS before being resuspended in 500 µL staining buffer (PBS containing 40 µg/ml propidium iodide, 100 µg/ml RnaseA, 0.1% Triton X-100), incubated on ice for ∼1hr and finally analyzed on flow cytometry.

### Identification of apoptosis related lncRNAs

The most variable genes related to apoptosis was identified using the approach presented by Brennecke et al^52^. A fit to the squared coefficients of variations against the means of normalized gene expressions (RPKM) was performed using the R function glmgam.fit(). The cell-to-cell variability of genes was ranked and the 75 apoptotic related genes (GO:0043065, positive regulation of apoptosis) with the greatest variability was used for PCA. Cell clusters were identified using the pam function of the R package ‘cluster’. Only cells assigned to the G1 cell cycle phase was considered for these analyses.

### qRTPCR

Total RNA was extracted using the Qiagen RNeasy mini kit (Qiagen cat no. 74106) followed by DNase treatment using the Ambion DNA-free DNA removal kit (Thermo Fisher Scientific, cat no. AM1906). Equal amounts of DNase treated RNA was used for preparing cDNA using SuperScript II (Thermo Fisher Scientific) and oligo(dT)_18_ primer according to the manufacturer’s recommendations. Quantification was carried out using Power SYBR Green (Thermo Fisher Scientific) on a StepOnePlus or ViiA7 Real-Time PCR system (Applied biosystems). The delta-delta Ct method was used for quantification of relative expression levels.

### Cloning / Generation of lentiviral U6 expressed shRNAs

Cloning of shRNAs into lentiviral vector was carried out as previously described^47^. Briefly, oligos were treated with T4 PNK (New England Biolabs), annealed and ligated into the Nhe1 and Pac1 restriction sites of the pHIV7-IMPDH2-U6 construct^47^.

### Lentiviral production

Lentiviral particles were produced as previously described^47,53^. Shortly, HEK293FT cells were transfected with pCHGP-2, pCVM-G pCMV-rev and pHIV7-IMPDH2-U6 at 1:0.5:0.25:1.5 ratio using SuperScript II and PLUS reagent (Thermo Fisher Scientific) in serum depleted DMEM media. Media was changed ∼6 hours post transfection to DMEM collection media (10% FBS (Thermo Fisher Scientific), 1% Penicillin/Streptomycin (Thermo Fisher Scientific), 1% Non-essential amino acids (Thermo Fisher Scientific), 1% Sodium-Pyruvate (Thermo Fisher Scientific), 0.1mM NEAA (Thermo Fisher Scientific) and 2mM L-Glut (Thermo Fisher Scientific), 0.37% Sodium Bicarbonate (Thermo Fisher Scientific) and 1x Viral Boost Reagent (Alstem Cell advancements). The viral supernatant was collected ∼48 hours post transfection, passed through a 0.45 µm filter (Sarstedt) and concentrated using PEG-it (SBI System Biosciences) according to the manufacturer’s recommendations.

### Generation of lentiviral transduced stable cell lines

NIH3T3 cells were transduced using low titer of lentiviral particles (< 10% of transduced cells). GFP+ cells were sorted at the CMB core facility at Karolinska Institutet.

### Colony formation assay

For stable NIH3T3 cell lines, cells were seeded at low density (500 cells / well) in 6 well plates. Media was changed every second to third day. After 10-14 days, cells were washed twice in PBS, fixed and stained for 20 minutes with 0.5% crystal violet, washed in H2O and finally let dry. Quantification was carried out in a two-step process by first manually counting the colonies and secondly measuring the crystal violet absorbance at 590nm by re-solubilize the stained cells in 10% acetic acid solution.

For siRNAs, NIH3T3 cells were seeded at 1000-2,500 cells / well in 6 well plates. Transfections were carried out 24 hours after seeding and the procedure as described above repeated.

### siRNA knockdown and mitomycin C treatment

NIH3T3 and primary cells were transfected using Lipofectamine RNAiMAX Transfection Reagent (Thermo Fisher Scientific) according to the manufacturer’s protocol. A final concentration of 10nM siRNA was used. MMC treatment was initiated 24 hours post transfections.

### Functional prediction of lncRNAs using allelic imbalance

Genes were first filtered for **1)** having at least 3 allelic read counts in 20 cells, **2)** not being imprinted (**Supplementary Table 1**) **3)** not being encoded on the X-chromosome and **4)** having one of the following Ensembl BioMart annotations; protein_coding (n=10,789), lncRNA (n=545), pseudogene (n=1), transcribed_processed_pseudogene (n=9), transcribed_unitary_pseudogene (n=4), unitary_pseudogene (n=1), transcribed_unprocessed_pseudogene (n=20), unprocessed_pseudogene (n=8).

Allelic imbalance of individual genes was measured as previously defined (CAST_allelicCounts_

/(CAST_allelicCounts_ + C57_allelicCounts_) - 0.5) and an allelic score (allelicImbalance_lncRNA_ + allelicImbalance_mRNA_ – diff(allelicImbalance_lncRNA_, allelicImbalance_mRNA_)) was calculated for each lncRNA:mRNA gene pair within 500 kb of the lncRNA-TSS. The allelic score of lncRNA:mRNA gene-pairs was compared to a permutation test where each lncRNA (n=542) was moved to 1,000 randomly selected mRNA gene positions (the 1,000 genomic loci were kept the same for all lncRNAs and required to have at least two other genes in proximity). The allelic score was computed for each lncRNA:mRNA gene-pair over the randomly selected genomic loci (within +/- 500k bp) and p-values finally calculated as: allelicScore_lncRNA:mRNA:random_ >= allelicScore_lncRNA:mRNA:real_ / n_randomGeneInteractions_.

### Functional prediction of lncRNAs using single-cell resolved RNA-expression

Coordinated allelic expression of lncRNA-mRNA gene pairs (on the single-cell level) was addressed for all lncRNA-mRNA gene pairs within +/- 500 kb of the lncRNA TSS (n=542 lncRNAs). The expression pattern for each gene pair (>= 3 allelic read counts) was first evaluated by a Fisher’s exact test (P_Real_, Benjamini-Hochberg adjusted). To estimate the background, each lncRNA was next translocated to 1,000 randomly selected gene locations and a Fisher’s exact test applied for all randomly generated gene pairs (P_Random_, Benjamini-Hochberg adjusted). lncRNA-mRNA gene pairs were considered significant if P_Real_ < 0.01 where P_Random_ < P_Real_ occurred in less than 1% of the permutated gene interactions.

### Estimation of RNA half-lives and decay rates

Primary adult mouse tail fibroblasts were treated with Actinomycin D (Sigma-Aldrich cat no. SBR00013-1ml) at a final concentration of 5 µG/ml in quadruplicates (one female CASTxC57 and one female C57Bl6, both in technical duplicates). RNA was extracted using the RNeasy mini kit (Qiagen cat no. 74106) at 0, 1, 2, 4, 7 and 10 hours of treatment and global levels of RNA measured by polyA^+^ RNA-seq. Briefly, ∼60 ng of DNase treated RNA (Ambion DNA-free cat no. AM1906) was prepared for sequencing using the Smart-seq2 protocol (slightly modified for bulk RNA-seq) and sequenced on an Illumina NextSeq-500 (High output kit v2.5, 75 cycles). Data was processed using the zUMIs v2.9.3e pipeline and genes filtered for having at least 10 read counts in all four samples in the untreated condition (t_0_). Using RPKMs, gene expression was first normalized to the untreated condition (setting t_0_ = 1) for each individual sample. To normalize the expression over Actinomycin D treated time points, we took advantage of previous estimates of RNA half-lives in mouse ES cells^31^. To normalize the data, we first identified a subset of control genes with half-life estimates 1h < t_1/2_ < 8h with at least 50 read counts at t_0_ in all four Actinomycin D treated samples. The expected expression level of the control genes was calculated (y = 1*exp(-k_control_*t)) and used to compute a ‘normalization factor’ (by taking the median) for each time point and sample, to which all genes were normalized to reach the final relative expression levels. Genes with shorter half-lives than 2h were excluded from the 7 and 10 hour time-points when calculating the ‘normalization factor’. To estimate half-lives, the normalized expression was fitted to an exponential decay curve (y = a*exp(-kx)) using the R package ‘drc’. Decay rates (λ) was finally calculated using the formula: t_1/2_ = ln(2) / λ. Genes with half-lives < 10 hours and burst duration < 72 hours were considered for downstream analysis.

### PI – AnnexinV staining

PI-AnnexinV staining was carried out using the Annexin-V-FLUOS Staining kit (Roche, cat no. 11858777001). Briefly, treated cells were harvested using Tryp-LE (including the cell media and PBS wash), washed once in PBS, and resuspended in 75-100 µL of Annexin-V labeling solution according to the manufacturer’s protocol. The samples were incubated for 15 minutes at room temperature (alternatively for 45 minutes on ice) followed by adding 400 µL of ice-cold Annexin-V incubation buffer. The samples were subsequently analyzed on a BD FACSMelody Cell Sorter.

### Ethical compliance

This study has been approved by the Swedish Board of Agriculture, Jordbruksverket: N343/12

