## Supplemental Figures 1-11 for "Transcriptional kinetics and molecular functions of long non-coding RNAs"

#### SUPPLEMENTARY FIGURES

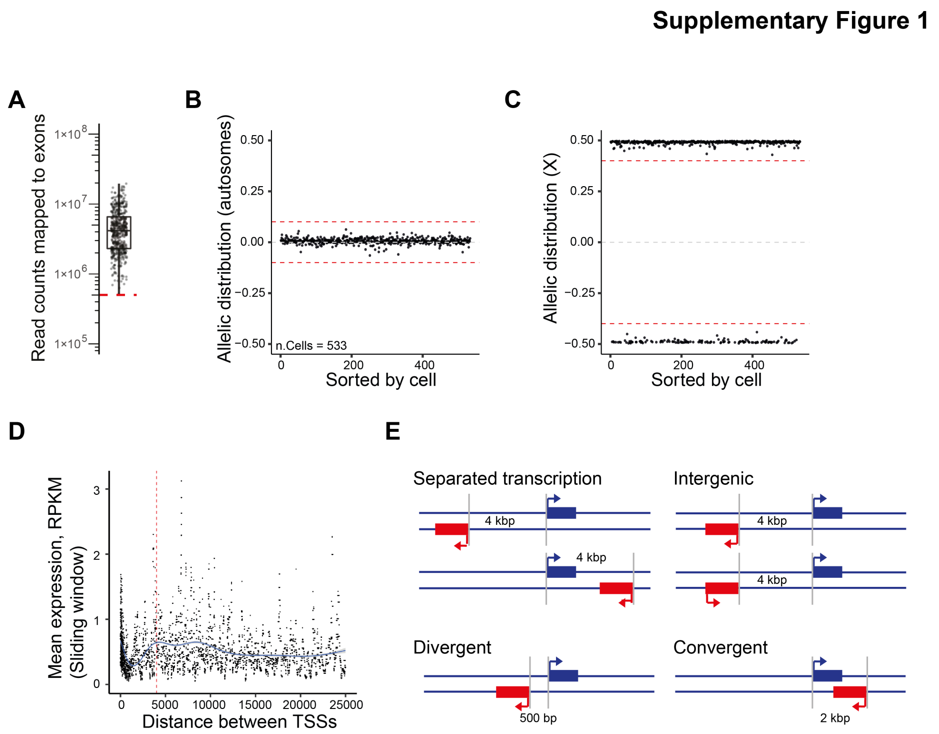


**Figure S1 | Quality controls for allele-sensitive scRNA-seq data.**

**(A)** Boxplot showing number of sequenced read counts mapped to genes (Smart-seq2 libraries, n=533 cells) (red line represents quality control cutoff). **(B)** Scatter plot showing the distribution of allele sensitive read counts (($counts_{CAST}$ / ($counts_{CAST}$ + $counts_{C57}$) - 0.5)) for non-imprinted autosomal genes (red dashed lines represent quality control cutoffs). **(C)** Scatter plot showing the distribution of allele sensitive read counts for non-escapee genes on the X-chromosome (red dashed lines represent quality control cutoff). **(D)** Mean expression of a sliding window (median RPKM, width=51) against distance between two promoters. The blue line denotes a generalized additive mode smoothing (gam) fitted to the sliding median. The red dashed line represents the cutoff in distance between two promoters for being considered as separated transcriptional unit (4 kb). **(E)** Schematic representation of gene organizations of genes and the nomenclature used in this manuscript.

**Supplementary Figure 2**

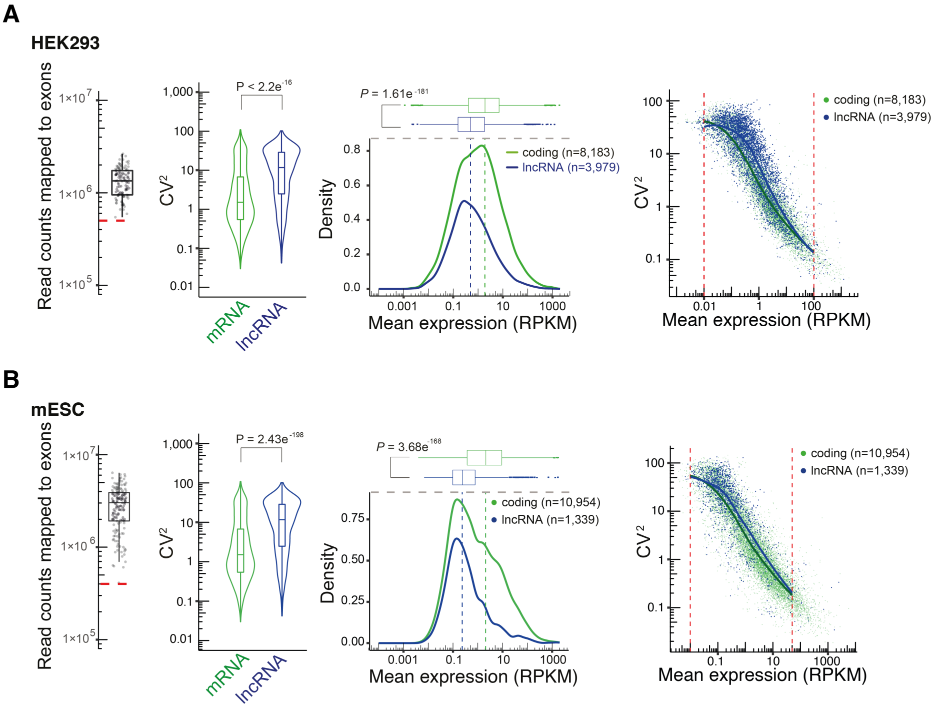


**Figure S2 | Complementary analyses of lncRNA expression patterns in HEK293 and mES cells.**

**(A-B)** Reproducing the analyses and results presented in Figure 1A,B,D,E using (A) HEK293 cell or (B) mouse embryonic stem cell data.

**Supplementary Figure 3**

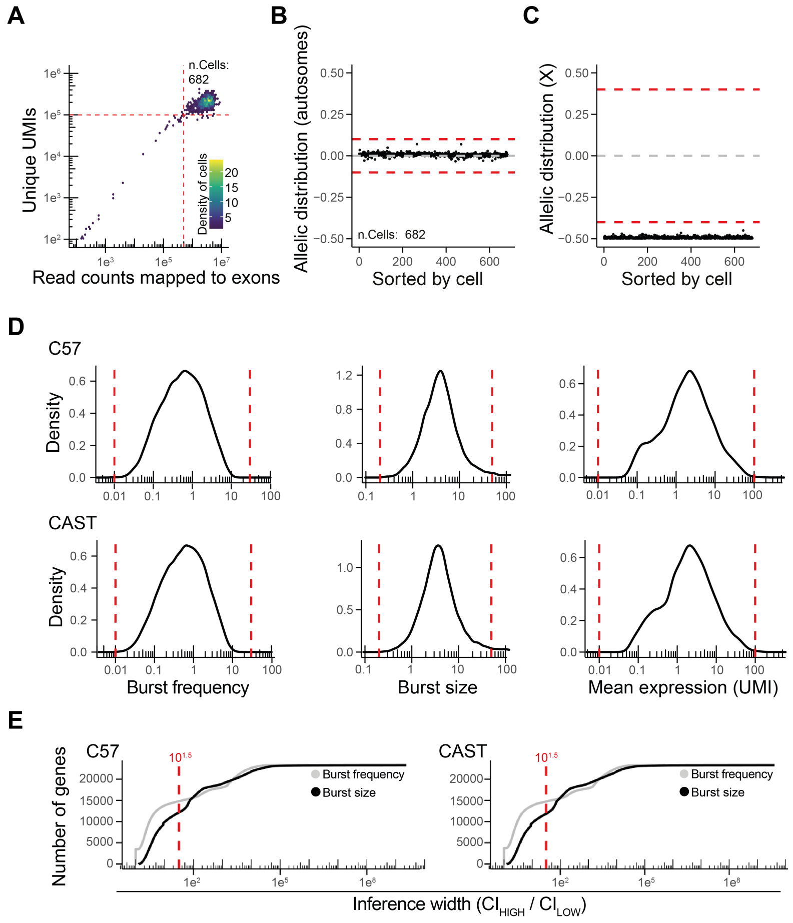


**Figure S3 | Quality controls and bursting inference data using Smart-seq3 data.**

**(A)** Scatter plot showing the number of sequenced read counts mapped to exons (Smart-seq3 libraries, n=682 cells passing quality control) (red lines represent quality control cutoffs). **(B)** Scatter plot showing the distribution of allele sensitive read counts for non-imprinted autosomal genes (red dashed lines represent quality control cutoffs). **(C)** Scatter plot showing the distribution of allele sensitive read counts for non-escapee genes on the X-chromosome (red dashed lines represent quality control cutoff). **(D)** Density plots for burst frequencies, burst sizes and mean expression (for allele distributed UMIs). Red dashed lines represent cutoffs for passing the quality control. **(E)** Scatter plots representing widths of confidence intervals (x-axis) against genes (y-axis, sorted by widths of confidence intervals) for burst frequencies and burst sizes. Red dashed lines represent cutoffs for passing the quality control.

**Supplementary Figure 4**

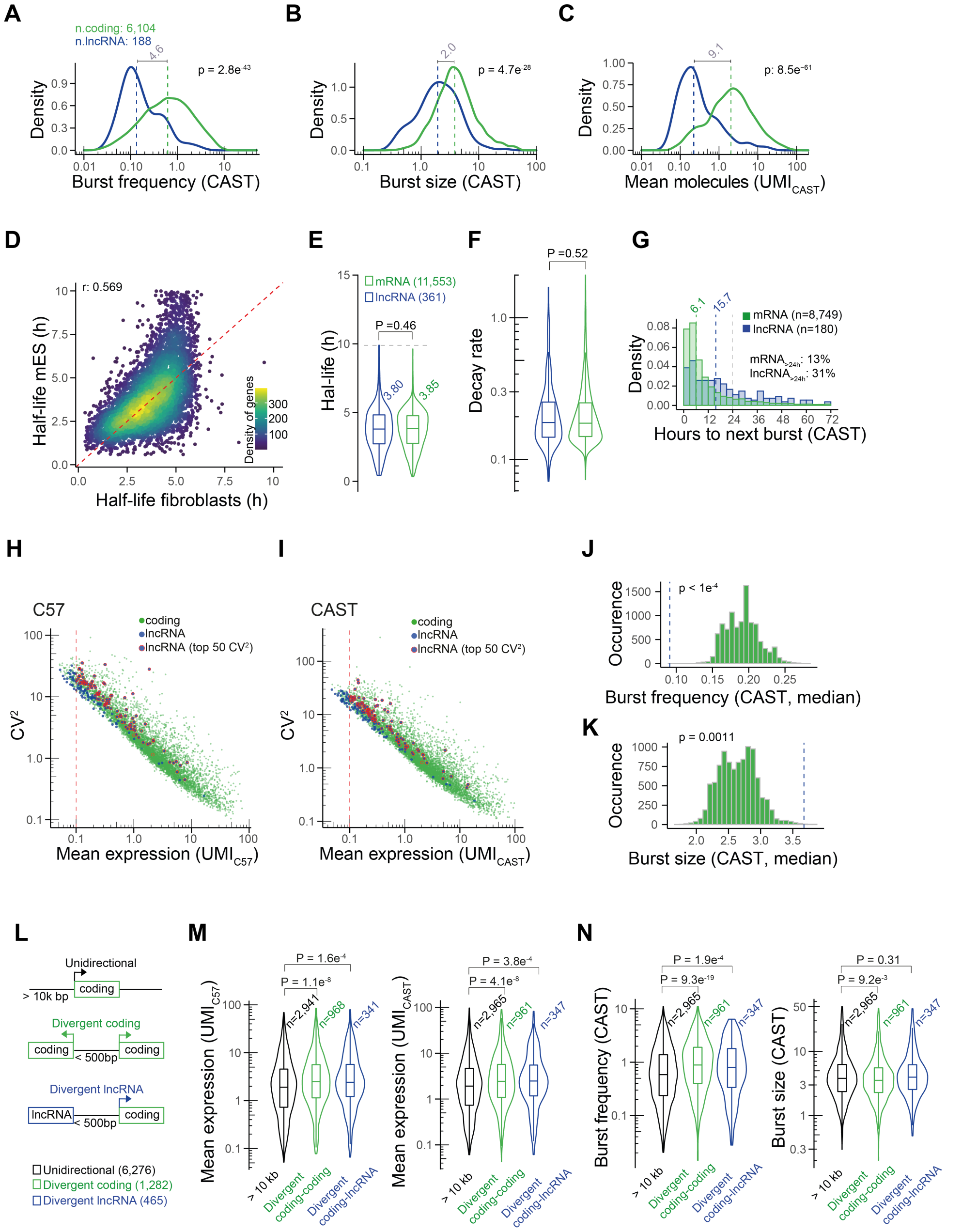


**Figure S4 | Transcriptional burst kinetics of lncRNAs and divergent promoters.**

**(A-C)** Density plots for (A) burst frequencies (B) burst sizes and (C) mean expression (for allele-distributed UMIs) for mRNAs and lncRNAs (showing the CAST allele). Dashed lines represent the median burst frequencies, sizes and mean expression for mRNAs and lncRNAs. The relative fold changes in burst frequencies, sizes and mean expression are annotated in grey. The p-values represent two-sided Wilcoxon rank-sum tests. **(D)** Scatter plot comparing estimated

RNA half-lives for mouse ES cells and mouse fibroblast cells. **(E-F)** Violin plots for (E) RNA half-lives and (F) RNA decay rates for mouse fibroblasts. P-values represent two-sided Wilcoxon rank-sum tests. **(G)** Histogram showing the duration between two bursts from the same allele (CAST), for mRNAs and lncRNAs. Dashed lines in green and blue represent the median duration between two bursts for mRNAs and lncRNAs, respectively. Dashed line in grey represents a duration of 24 hours between two bursts. **(H-I)** Scatter plot of mean expression (for allele-distributed UMIs) against the $CV^2$ for lncRNAs (blue) and mRNAs (green) for the (H) C57 and (I) CAST genomes. The $CV^2$ of each lncRNA was ranked to 100 mRNAs of similar mean expression and the top 50 ranked lncRNAs are highlighted in red. The red dotted line denotes the lower expression cutoff for lncRNAs to be included in the analysis. **(J-K)** Histograms showing median (J) burst frequencies (K) and burst sizes for sampled expression-matched sets of mRNAs. Each lncRNA (n=50, identified in Figure S4I) was matched with 10 mRNAs of similar expression followed by subsampling one expression-matched mRNA for each lncRNA. The P-value represent the outcome of the permutation test, where the observed burst parameters (lncRNAs, median) was higher (for burst frequencies) or lower (for burst sizes) than the burst parameters (median) obtained from 10,000 permutations. **(L)** Schematic representation of divergent and unidirectional transcribing promoters. **(M)** Violin plots for mean expression of mRNAs (allele-distributed UMIs) for divergent and unidirectional promoters for the C57 and CAST alleles. The P-values represent a two-sided Wilcoxon rank-sum test. **(N)** Violin plots for unidirectional and divergent promoters representing burst frequencies and burst sizes, for the CAST allele. P-values represent two-sided Wilcoxon rank-sum test.

**Supplementary Figure 5**

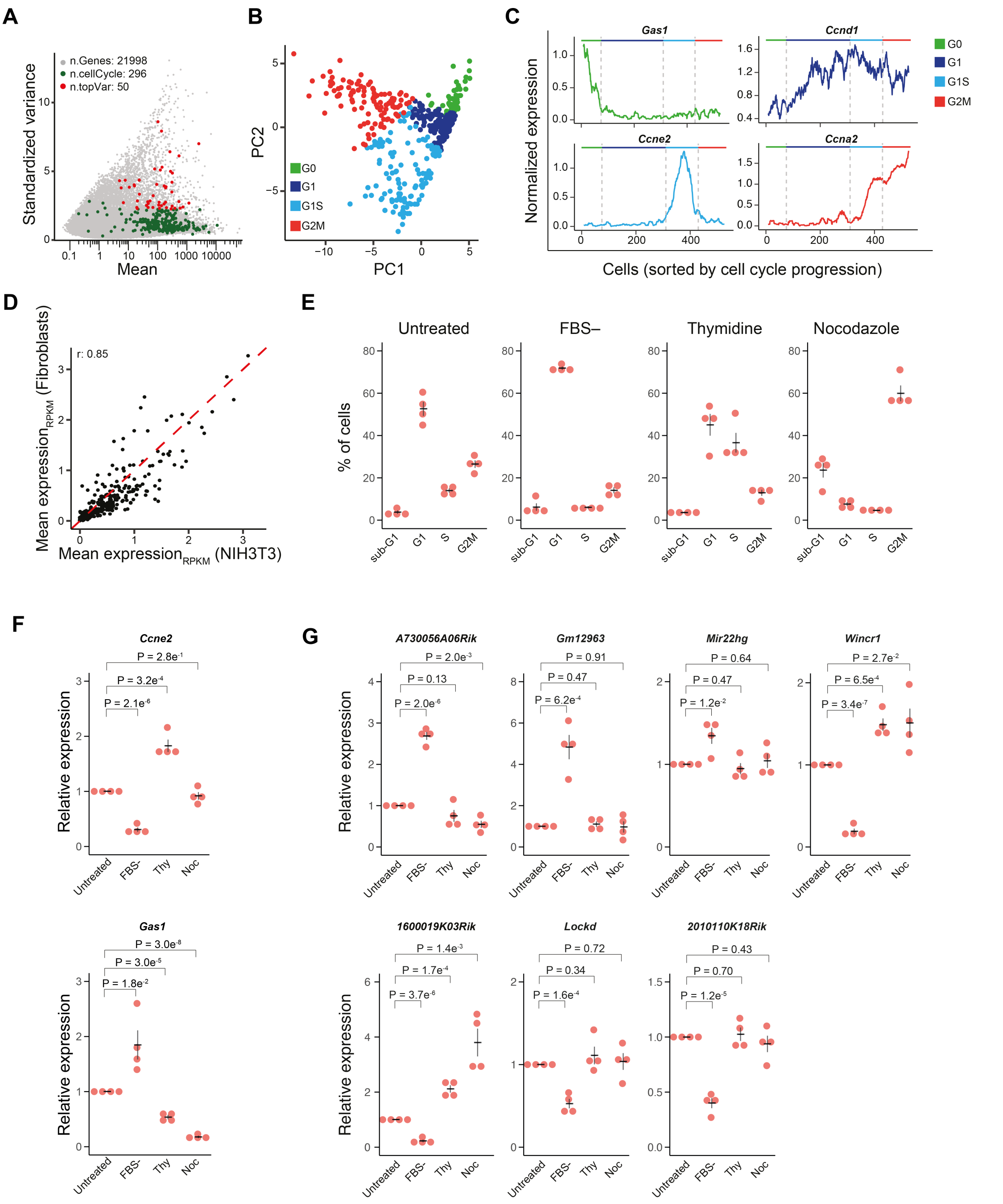


#### Figure S5 | Detailed analysis of cell cycle associated lncRNAs in the scRNA-seq data.

**(A)** Scatter plot highlighting the 50 most variable cell cycle genes[36] identified by using the R package Seurat[35], on top of a scatter plot showing mean and standardized variance in expression over cells. **(B)** Low dimensional PCA projections of cells based on the most variable genes identified in (A). Cells are colored according to cell cycle annotations in Figure 3A. **(C)**

Expression pattern of representative marker genes across cells, ordered according to cell cycle progression. The expression represents a sliding window (width = 15) of the mean expression (scale-factor and log10 normalized by the R package Seurat). **(D)** Scatter plot representing the mean expression of cell cycle genes in mouse primary fibroblasts (n=533 cells) against lentiviral transduced (shRNA-Control) NIH3T3 (n=147 cells) (scale-factor and log10 normalized using the R package Seurat, r represents the Spearman correlation). **(E)** Barplots showing cell cycle distribution of NIH3T3 cells upon various cell cycle synchronizations by indicated compounds. **(F)** Relative expression measured by qRTPCR of two cell cycle marker genes upon cell cycle synchronization in NIH3T3 cells. Samples were standardized to RNA input. **(G)** Relative expression measured by qRTPCR for candidate lncRNAs upon cell cycle synchronization in NIH3T3 cells. Samples were standardized to RNA input. (Treatments in Figure S5E-G represent; FBS- (serum starvation 0.1%), Thy (Thymidine), Noc (Nocodazole))

**Supplementary Figure 6**

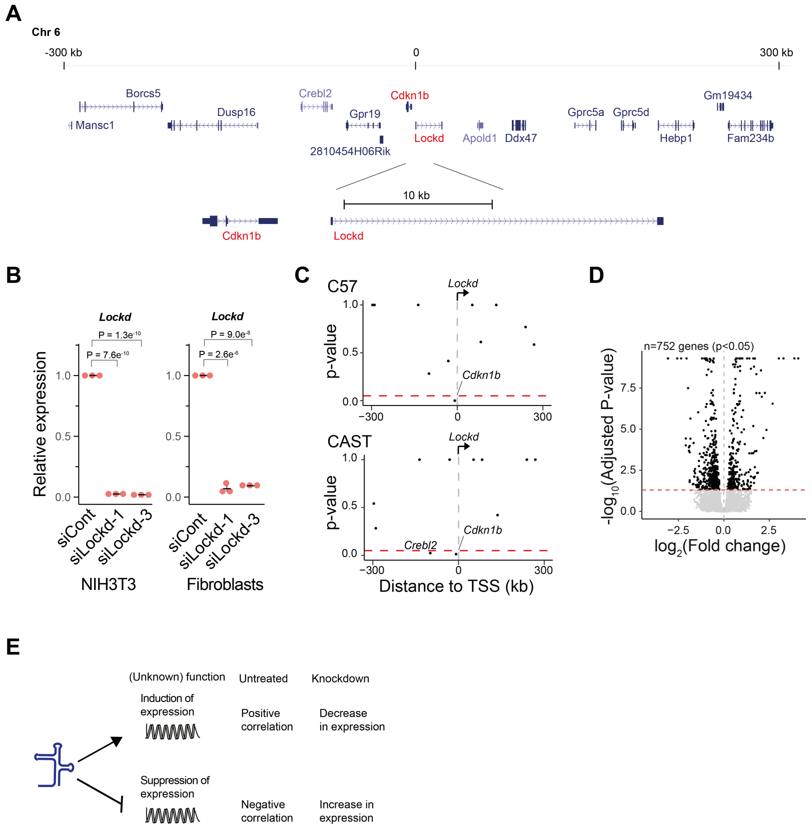


**Figure S6 | Details of the *Lockd* harboring genomic loci and experiments.**

**(A)** A modified schematic from the UCSC genome browser representing the *Cdkn1b-Lockd* gene locus. **(B)** Relative expression of *Lockd* upon siRNA induced knockdown in NIH3T3 cells (n=4) and primary fibroblasts (n=4) measured by qRTPCR, and in control cells. P-values represent a two-sided student's t-test, error bars represent the s.e.m. **(C)** Scatter plot showing p-values for co-expression of *Lockd* (Fisher's exact test, gene considered expressed if supported by >= 3 allelic counts) against genes in proximity (+/- 300 kb of TSS-*Lockd*) for the CAST and C57 genomes. The red dashed lines denote threshold for significance (p<0.05). **(D)** Scatter plot representing p-values (analyzed by SCDE) against fold changes of shLockd treated NIH3T3 cells. The red dashed line denotes threshold for significance (p<0.05). **(E)** Schematic representation for the intersection of fold change (shLockd / shControl) against gene-gene correlations in shControl treated cells.

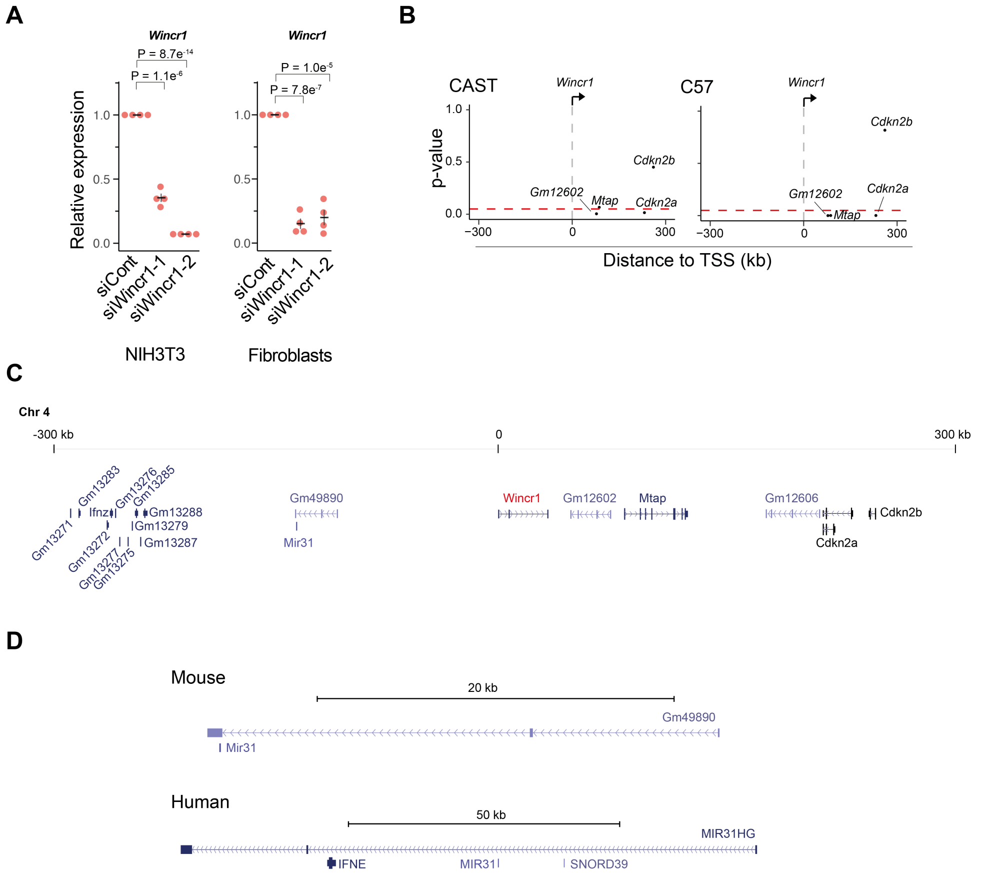


**Figure S7 | Details of the *Wincr1* genomic loci and experiments.**

**(A)** Relative expression of *Wincr1* upon siRNA induced knockdown in NIH3T3 cells measured by qRTPCR (n=5, p-values represent a two-sided student's t-test, error bars represent the s.e.m.), and in control cells. **(B)** Scatter plots showing p-values for co-expression of *Wincr1* (Fisher's exact test, gene considered expressed if supported by >= 3 allelic counts) against other genes in proximity (+/- 300 kb of TSS-*Wincr1*) for the CAST and C57 genomes, respectively. The red dashed lines denote threshold for significance (p<0.05). **(C)** A schematic from the UCSC genome browser representing the *Wincr1* gene locus. **(D)** A schematic from the UCSC genome browser comparing the genomic loci of the microRNA-31 host gene in human and mouse.

**Supplementary Figure 8**

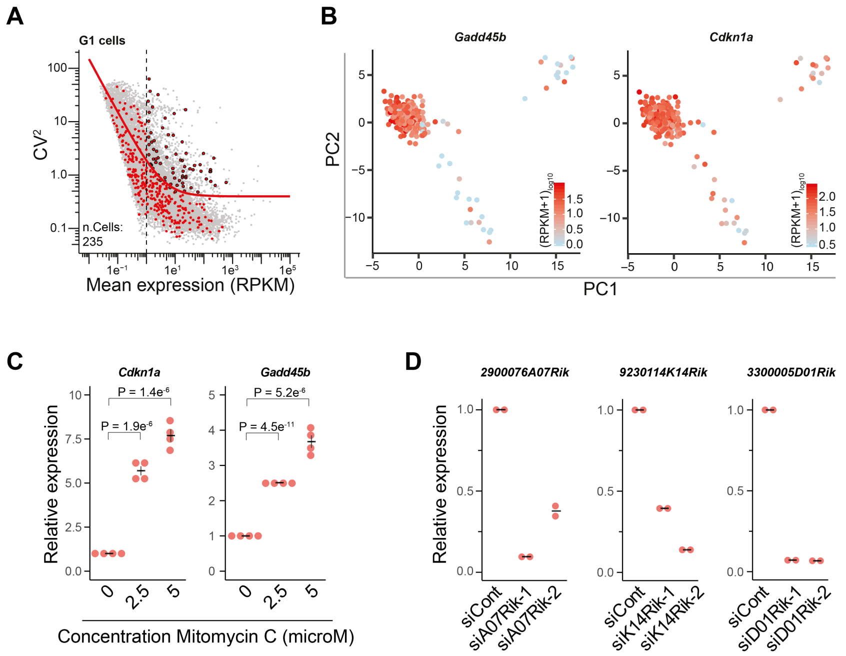


**Figure S8 | Detailed analyses of apoptosis signals in the scRNA-seq data.**

**(A)** Scatter plot showing the 75 most variable genes related to apoptosis identified using the method presented by Brennecke et al[52], on top of a scatter plot showing mean expression levels (x-axis) and $CV^2$ (y-axis). **(B)** Low dimensional PCA projections of cells based on the most variable genes identified in (A), colored by the expression of *Gadd45b* and *Cdkn1a*. **(C)** Relative expression measured by qRTPCR for *Gadd45b* and *Cdkn1a* in NIH3T3 cells treated with MMC (n=4, p-values represents a two-sided student's t-test, error bars represent the s.e.m.). **(D)** Relative expression of candidate lncRNAs upon siRNA induced knockdown in NIH3T3 cells measured by qRTPCR (n=2).

Supplementary Figure 9

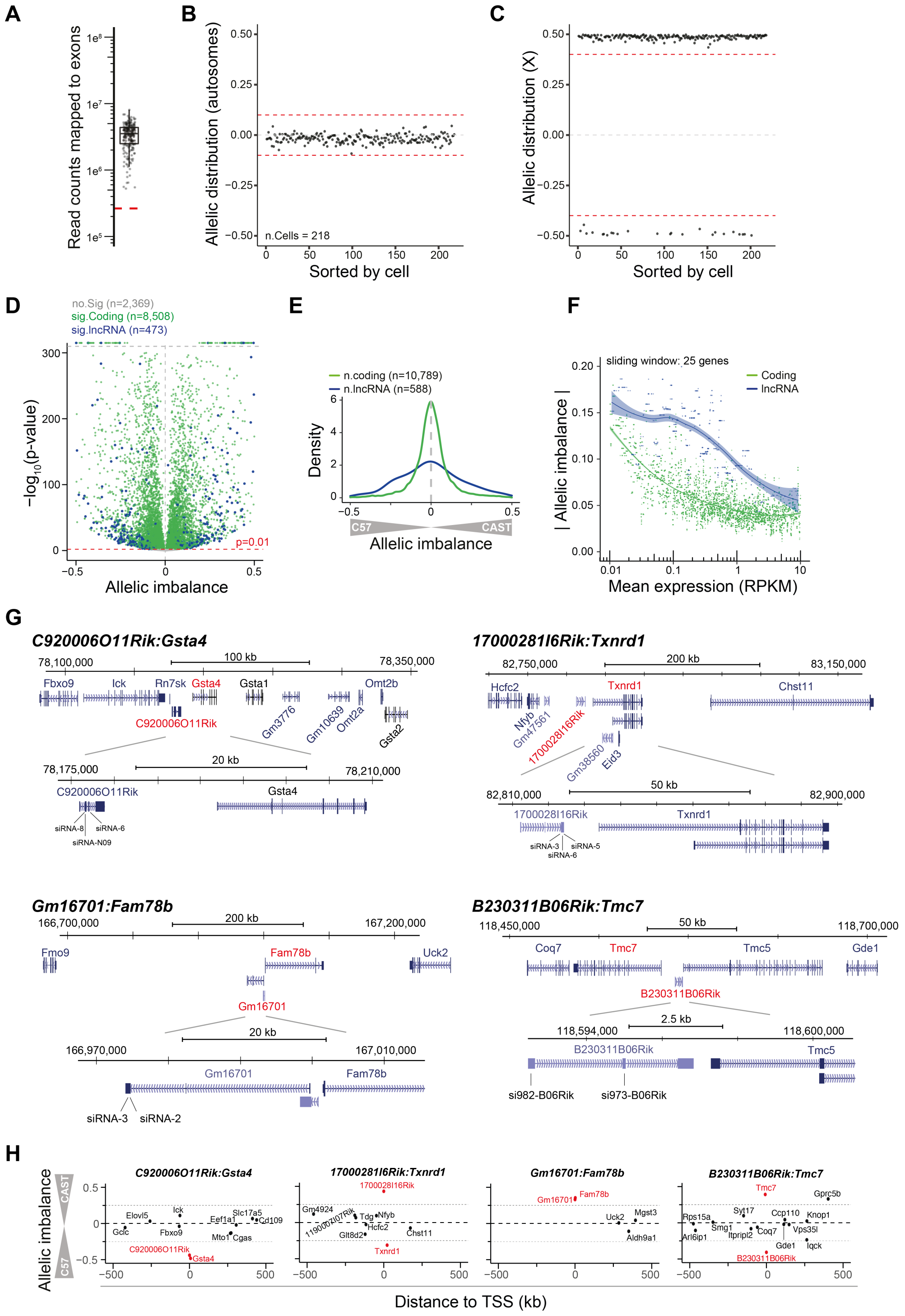

### Figure S9 | Control experiments on allelic imbalance in fibroblast cells

**(A-C)** Quality control for Smart-seq2 libraries from CASTxC57 primary fibroblast cells. **(A)** Boxplot showing number of sequenced reads counts mapped to exons (Smart-seq2 libraries, n=218 cells) (red line represents quality control cutoff). **(B)** Scatter plot showing the distribution of allele sensitive read counts for non-imprinted autosomal genes (red dashed lines represent quality control cutoffs). **(C)** Scatter plot showing the distribution of allele sensitive read counts for non-escapee genes on the X-chromosome (red dashed lines represent quality control cutoff). **(D)** Scatter plot of the allelic bias against p-values (binomial test, Benjamini-Hochberg adjusted) across fibroblasts (n=751). **(E)** Density plot summarizing observed allelic imbalance of lncRNAs and mRNAs across fibroblasts (n=751). **(F)** Mean expression towards the median of allelic imbalance of a sliding window (width = 25) for mRNAs and lncRNAs. The green and blue lines denote a loess fit to the sliding window. **(G)** A modified schematic from the UCSC genome browser representing the gene loci of four candidate lncRNA-mRNA gene pair interactions. **(H)** Scatter plot representing allelic bias of genes within +/-500kb (of lncRNA TSS) of candidate genes. Candidate lncRNA-mRNA gene pair interactions are colored in red.

**Supplementary Figure 10**

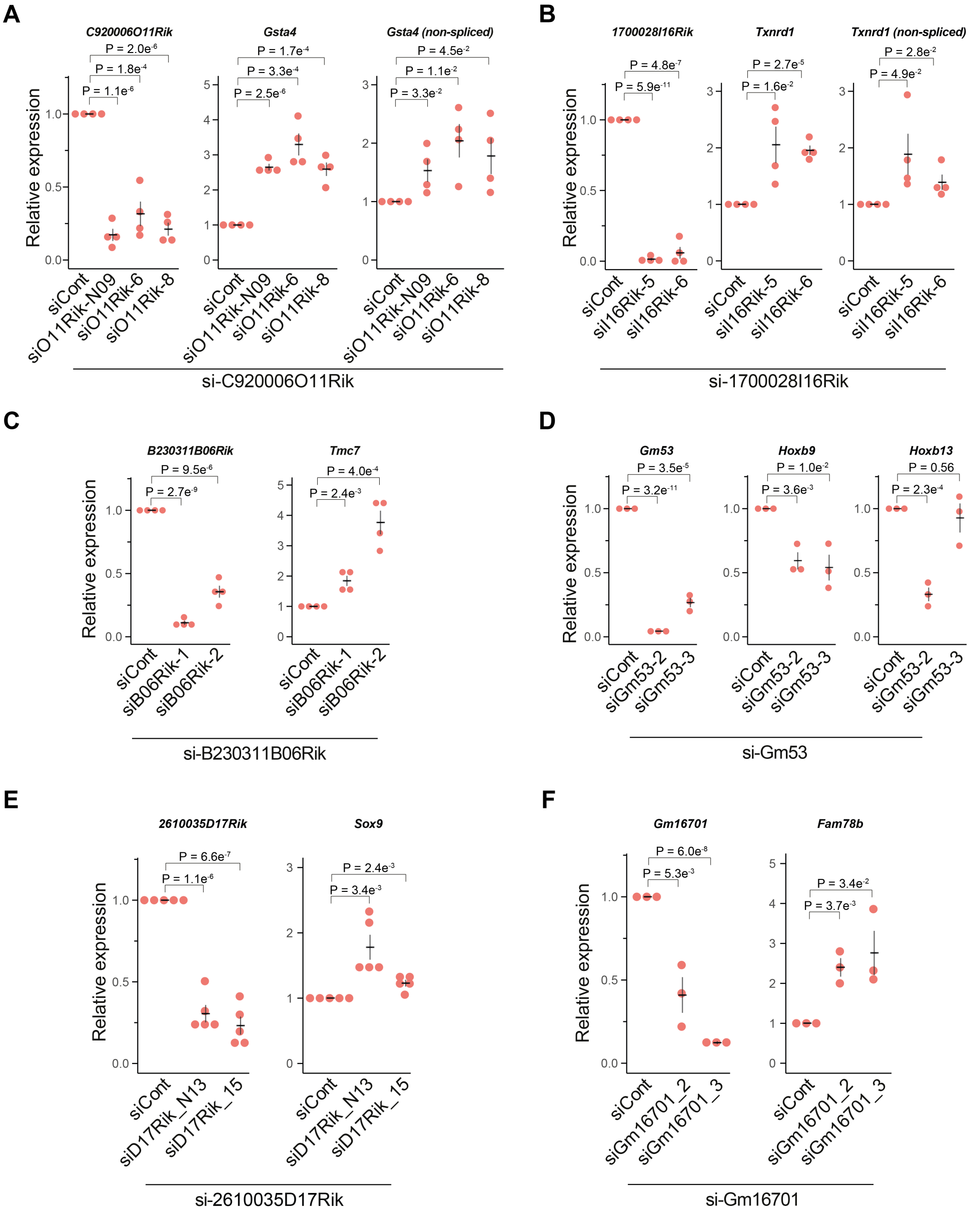


**Figure S10 | Validation experiments on lncRNA-mRNA interactions in fibroblast cells.**

**(A)** Relative expression levels of spliced and unspliced transcripts of *Gsta4* upon siRNA induced knockdown of *C920006O11Rik* measured by qRTPCR (n=4), **(B)** Relative expression levels of spliced and unspliced transcripts of *Txnrd1* upon siRNA induced knockdown of *1700028I16Rik* measured by qRTPCR (n=4). **(C)** Relative expression levels of spliced transcripts of *Tmc7* upon siRNA induced knockdown of *B230311B06Rik* measured by qRTPCR (n=4). **(D)** Relative expression levels of spliced transcripts of *Hoxb9* and *Hoxb13* upon siRNA induced knockdown of *Gm53* measured by qRTPCR (n=4) **(E)** Relative expression levels of spliced transcripts of *Sox9* upon siRNA induced knockdown of *2610035D17Rik* measured by qRTPCR (n=5). **(F)** Relative expression levels of spliced transcripts of *Fam78b* upon siRNA induced knockdown of *Gm16701*measured by qRTPCR (n=4). (A-F) P-values represent a two-sided student's t-test, error bars represent the s.e.m.

**Supplementary Figure 11**

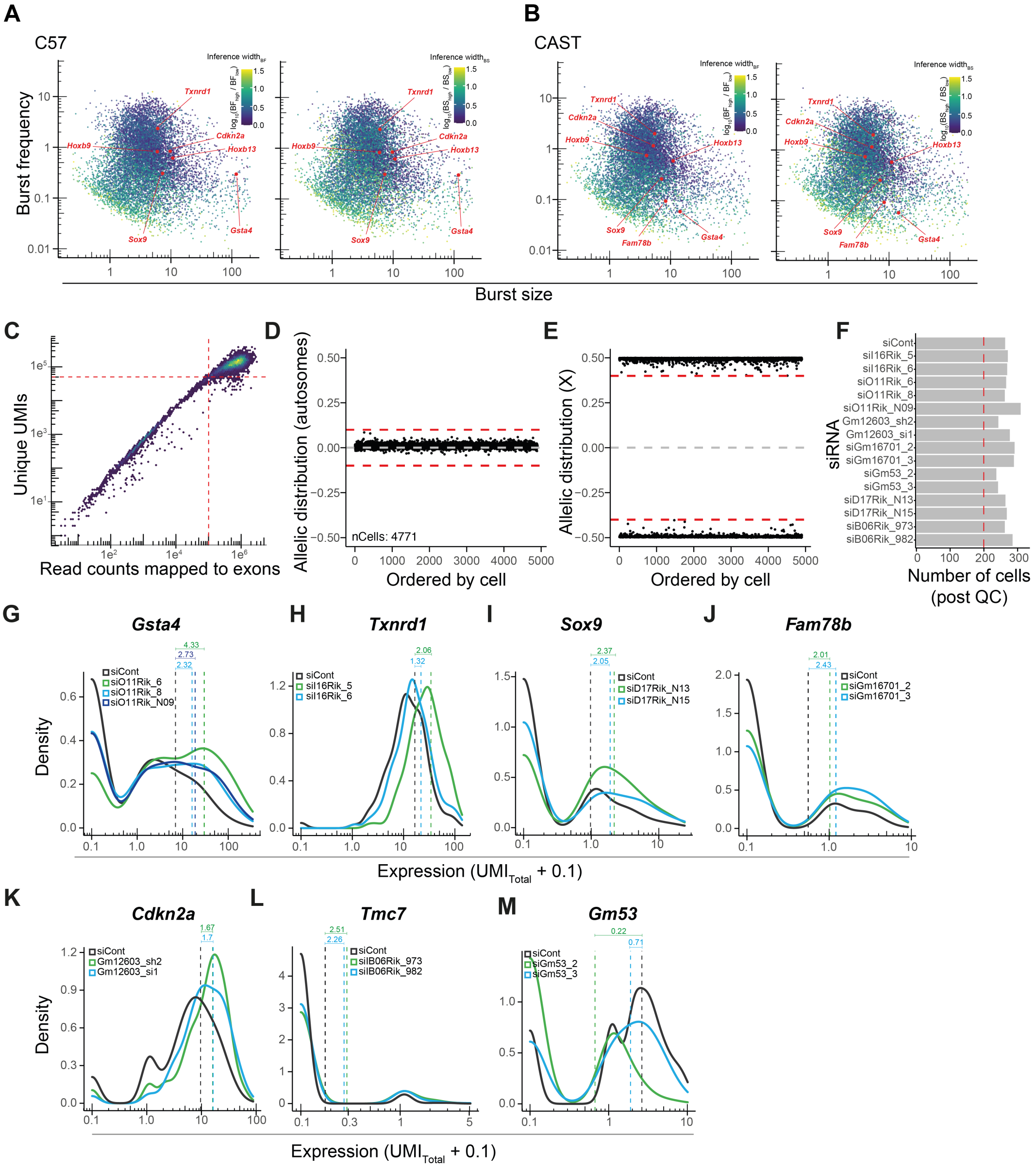

**Figure S11 | Detailed analyses of lncRNAs' effect on burst kinetics of nearby mRNAs.**

**(A-B)** Scatter plots showing the burst size against the burst frequency for the (A) C57 and (B) CAST genomes. Genes are colored according to the width of a 95% confidence interval ($CI_{high}/CI_{low}$). **(C-E)** Quality control for Smart-seq3 libraries from siRNA treated primary fibroblast cells. **(C)** Scatter plot showing the number of unique UMI reads against the number of read counts mapped to exons (red lines represent quality control cutoffs). **(D)** Scatter plot showing the distribution of allele sensitive read counts for non-imprinted autosomal genes (red dashed lines represent quality control cutoffs). **(E)** Scatter plot showing the distribution of allele sensitive read counts for non-escapee genes on the X-chromosome (red dashed lines represent quality control cutoffs). **(F)** Barplot representing the number of cells passing quality control for individual siRNA treatments. **(G-L)** Density plots representing mean expression (UMIs) of modulated genes upon siRNA induced suppression of lncRNAs for (G) *Gsta4*, (H) *Txnrd1*, (I) *Sox9*, (J) *Fam78b*, (K) *Cdkn2a* and (L) *Tmc7*. (G-L) Mean expression fold changes for individual siRNAs are quantified and highlighted in green and blue. Vertical dashed lines represent mean expression levels. **(M)** Density plot representing expression of the lncRNA Gm53 upon siRNA induced knockdown. Vertical dashed lines represent mean expression levels.
